# Climatic niche change of fish is faster at high latitude and in marine environments

**DOI:** 10.1101/853374

**Authors:** Luana Bourgeaud, Jonathan Rolland, Juan David Carvajal-Quintero, Céline Jézéquel, Pablo A. Tedesco, Jérôme Murienne, Gaël Grenouillet

## Abstract

Change in species’ climatic niches is a key mechanism influencing species distribution patterns. The question of which factors impact niche change remains a highly debated topic in evolutionary biology. Previous studies have proposed that rates of climatic niche change might be correlated with climatic oscillations at high latitude or adaptation to new environmental conditions. Yet, very few studies have asked if those factors are also predominant in aquatic environments. Here, we reconstruct the climatic niche changes of fish species on a new phylogeny encompassing 12,616 species. We first confirm that the rate of niche change is faster at high latitude and show that this association is steeper for freshwater than for marine species. We also show that freshwater species have slower rates of niche change than marine species. These results may be explained by the fact that freshwater species have larger climatic niche breadth and thermal safety margin than marine species at high latitude. Overall, our study sheds a new light on the environmental conditions and species features impacting rates of climatic niche change in aquatic habitats.

---

Species response to different environmental conditions across geological time is an essential process shaping global macro-ecological patterns of species distribution^1–5^. The increasing availability of large scale phylogenetic and distributional data has allowed for the explore the temporal dynamic of species ecological niches by calculating rates of niche change (or niche shifts) through time^6–9^. Understanding the factors affecting rates of niche change remains an open question at the crossroad between ecology and evolution^10^. Previous studies in terrestrial environments have shown a potential role of endothermy^11^, life-history strategy^12, 13^, niche breadth^5, 14^ and latitude^15^. Interestingly, it is still unknown whether those factors are ubiquitous and drive niche change in other environments than terrestrial. We propose to study the relationship between latitude and rates of climatic niche change in aquatic species (marine and freshwater) and explore how inhabiting different environments affects rates of climatic niche change. For this, we studied ray-finned fish (Actinopterygii) given that they occur in both freshwater and marine environments with similar diversity levels^16^, and that deep phylogenetic relationships for fish are well resolved^17^.

Recent work on terrestrial endotherms has proposed that rates of climatic niche change might be higher in the temperate regions than in the tropics^7, 15, 18^. Indeed, higher rate of niche change might be related to an increase of the strength of divergent selection at high latitude^15^. For example, extinction related to climatic oscillations^19^ should promote ecological opportunity at high latitude^20, 21^ whereas competition related to high species richness in the tropics may impede change in distributions and niche change^22^.

Besides latitude, we also expect that differences between marine and freshwaters environments^23^ would potentially impact rates of climatic niche change. We can formulate two competing hypotheses concerning the rates of change between freshwater and marine environments. First, the rate of climatic niche change might be faster in freshwaters than in marine habitats, because freshwaters are more spatially fragmented^24^, which would potentially facilitate speciation^25, 26^ and niche divergence related to the colonization of isolated rivers and lakes. Second, climatic niche change might be faster in marine lineages which live in larger volumes of water with larger distributional ranges^27^ and can potentially have access to a higher number of niches^28^ leading to more opportunities for niche change. Combining a new phylogeny of more than 12,000 species of fish with a large dataset of distribution and climatic data, we explore the differences in rates of climatic niche change across latitude in marine and freshwater environments.

We compiled a dataset of 6,104,127 georeferenced occurrences for 6,627 species of fish (3,622 marine, 3,005 freshwater species) across the world. For each species, we calculated the average of the maximum water temperature of warmest month (T_max_) and the minimum water temperature of coldest month (T_min_) for all the occurrences as an estimation of species current climatic niche (see Methods). We also reconstructed a dated phylogeny for 12,616 fish species (40% of the 28,900 described species) combining mitochondrial genetic data with a backbone topology built from the phylogenetic classification of bony fish^17^ (version v4, see Methods and Supplementary Methods). Independently for each fish family (aged between 4.8 and 139.48 Myr), we modeled temperature change along the branches of the phylogeny as a Brownian motion process and reconstructed ancestral values. From the reconstructed values, we computed species rates of absolute temperature (T_max_ and T_min_) niche change to investigate the intensity of change of species niche position (see Methods).

Following this approach, we found that rates of absolute temperature niche change were lower at lower latitudes (Fig. 1, Supplementary Table 1). Using Bayesian mixed-effect models, which account for phylogenetic relatedness, the age of the family and the species sampling, we found a positive quadratic effect of latitude on rates of T_max_ and T_min_ change (Fig. 2, Supplementary Table 2). To illustrate the geographical pattern of rate of temperature niche change, we mapped mean rates of temperature niche change for the species community of each 5° grid cell of marine environment and each freshwater drainage basin. We revealed a tropical belt of low rates of absolute temperature niche change while rates were higher toward the poles in particular in the northern hemisphere (median rate of T_max_ change in the tropics (between −23.4 and 23.4°): 0.043 [Interquartile range: 0.018, 0.103] °C.Myr^-1^ against 0.100 [0.044, 0.250] °C.Myr^-1^ in the temperate region and T_min_ in the tropics 0.089 [0.050, 0.171] °C.Myr^-1^ against 0.111 [0.055, 0.229] °C.Myr^-1^ in the temperate region) (Fig. 1). We also investigated the direction of the niche change to identify whether niche change was toward hotter or colder temperatures (Supplementary Fig. 1, Supplementary Fig. 2). In addition to show higher rates of change for species climatic niche position toward the pole, we show that rates of absolute niche breadth change are faster in the temperate region meaning that niche breadth varied more through time in this region than in the tropics (Supplementary Fig. 3).

**Fig. 1:**
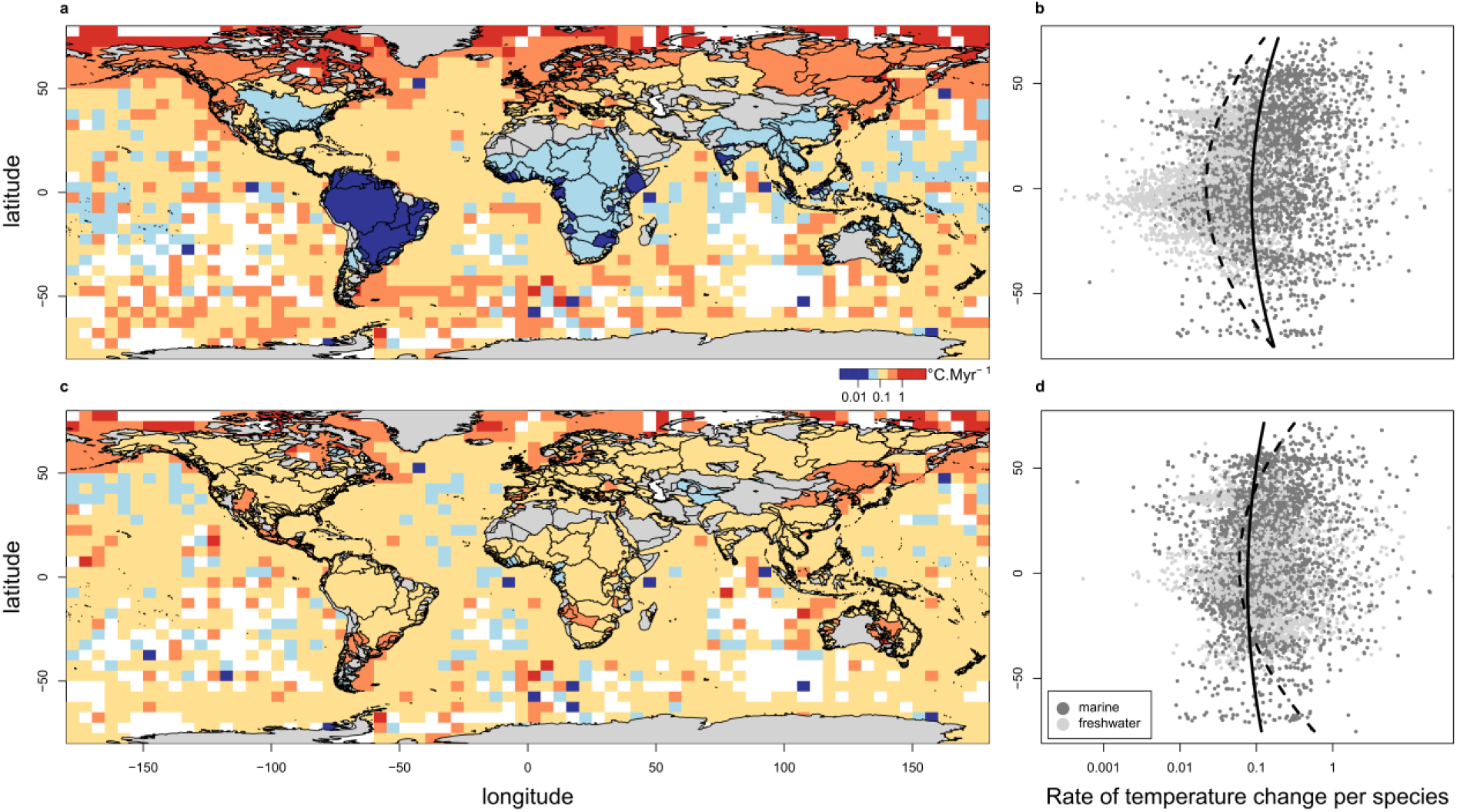
Geographical pattern of rates of absolute temperature change. Distribution of (a) rates of maximum temperature (T_max_) and (c) minimum temperature (T_min_) change across the globe (mean rate for the species community in each grid cell for marine species and mean rate for the fish community in each drainage basin for freshwater species). Absence of data is shown in grey on land and in white in the oceans. Relationship between rate of (b) T_max_ and (d) T_min_ change and latitude. Curves are built using the median estimate of the effect of latitude on rates of temperature change obtained using bayesian mixed-effect models accounting for phylogenetic relatedness between species (Supplementary Table S6).

**Fig. 2:**
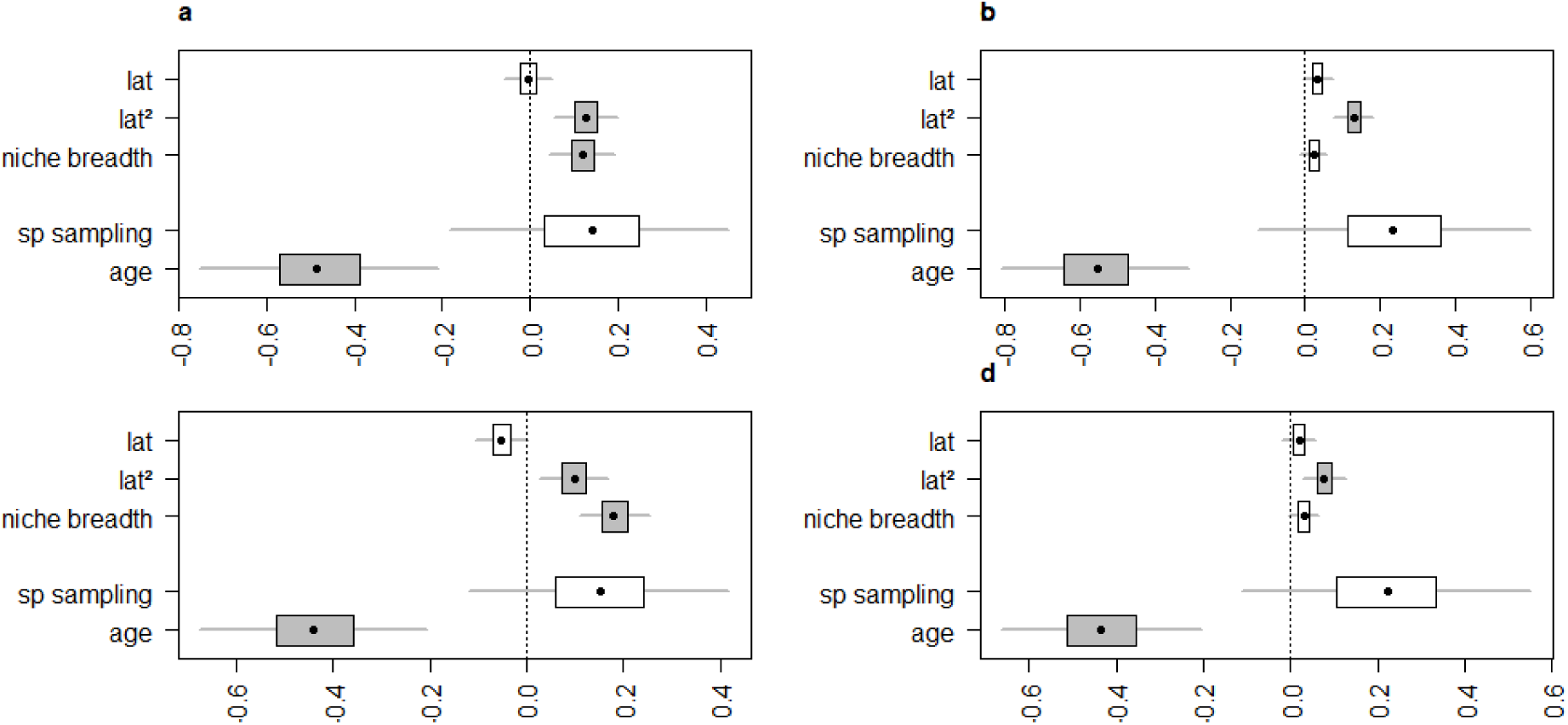
Effect of latitude, niche breadth, species sampling and clade age on rates of absolute T_max_ (a, b) and T_min_ (c, d) change for freshwater (a, c) or marine species (b, d). We used Bayesian mixed-effect models (MCMCglmm) to account for phylogenetic relatedness between species. Variables were scaled to obtain comparable estimates. The species sampling (sp sampling) is the number of species in the family analysed. Models were run on the 100 phylogenies using the R package Multree^57^. Boxplots represent the 95 per cent highest posterior density. Boxplots colored in grey show significant effects as they do not overlap zero. See Supplementary Table 2 for detailed results.

Several reasons can explain why rates of temperature change are higher in the temperate region^15^. First, climate heterogeneity in the temperate zone may be one of the factors driving climatic niche change^29^. Climate seasonality is much stronger in the temperate region as reflected by the larger niche breadth of temperate species (Supplementary Fig. 4). At a larger temporal scale, paleo-climatic fluctuations have been more intense in the temperate region^30, 31^ while the tropical climate has been much more stable. Both levels of climatic heterogeneity may have increased the opportunity for climatic niche change resulting in higher rates of temperature change for temperate species. Second, higher species richness in the tropics may prevent species geographic expansion (through competitive exclusion) and thus reduce the opportunity for climatic niche change^22^. On the contrary, species richness is lower in the temperate zone partially due to high levels of extinction as a result of the rapid changes in ecological conditions which might facilitate ecological opportunity and climatic niche change^32, 33^. Low species diversity and high ecological opportunity have also been proposed as an explanation for the higher rates of recent speciation in the temperate region^21^. Specifically, Rabosky et al.^34^ showed that current speciation rates for marine fish are faster in the temperate zone. Although we cannot demonstrate a causal effect of rates of climatic niche change on speciation with our data, we provide a new piece of evidence suggesting that speciation in fish could be related to species ability to adapt to new climatic conditions^3, 5, 35^.

Although we found a latitudinal gradient in the rates of niche change for both marine and freshwater species, we showed that rates of temperature change were different between the two habitats. For a given latitude, we found that rates of T_max_ and T_min_ change, were lower for freshwater species (T_max_ median rate: 0.033 [Interquartile range: 0.013, 0.074] °C.Myr^-1^ and T_min_ median rate: 0.090 [0.045 0.176] °C.Myr^-1^) than for marine species (T_max_: 0.100°C.Myr^-1^ [0.044 0.245] °C.Myr^-1^ and T_min_: 0.103 [0.057 0.210] °C.Myr^-1^) (Fig. 3). Phylogenetic generalized least square (pgls) regressions accounting for phylogenetic relatedness and phylogenetic uncertainty confirmed that rates of T_max_ change were lower for freshwater species (Fig. 3, Supplementary Table 3). The result was similar when we separated freshwater species from brackish water species (Supplementary Fig. 5, Supplementary Table 4). Higher rates of temperature change in marine environment were also found at the family level (Appendix 1 and Supplementary Fig. 6). We suggest that this result might be related to the differences in habitat heterogeneity and dispersal constraints between marine and freshwater habitats. Marine species tend to occupy a well-connected environment^28^ with a larger range of depths. They also have high dispersal capacities and large population sizes across wide ranges^36, 37^, which may lead them to be exposed to a wider range of climatic conditions resulting in more opportunities for climatic niche change. Moreover, modes of speciation might influence rates of climatic niche change. As freshwater habitats are more spatially fragmented^24^, they are thought to provide more opportunities for allopatric speciation without niche change^26^ whereas oceans might provide greater opportunities for speciation along ecological boundaries^38^.

**Fig. 3:**
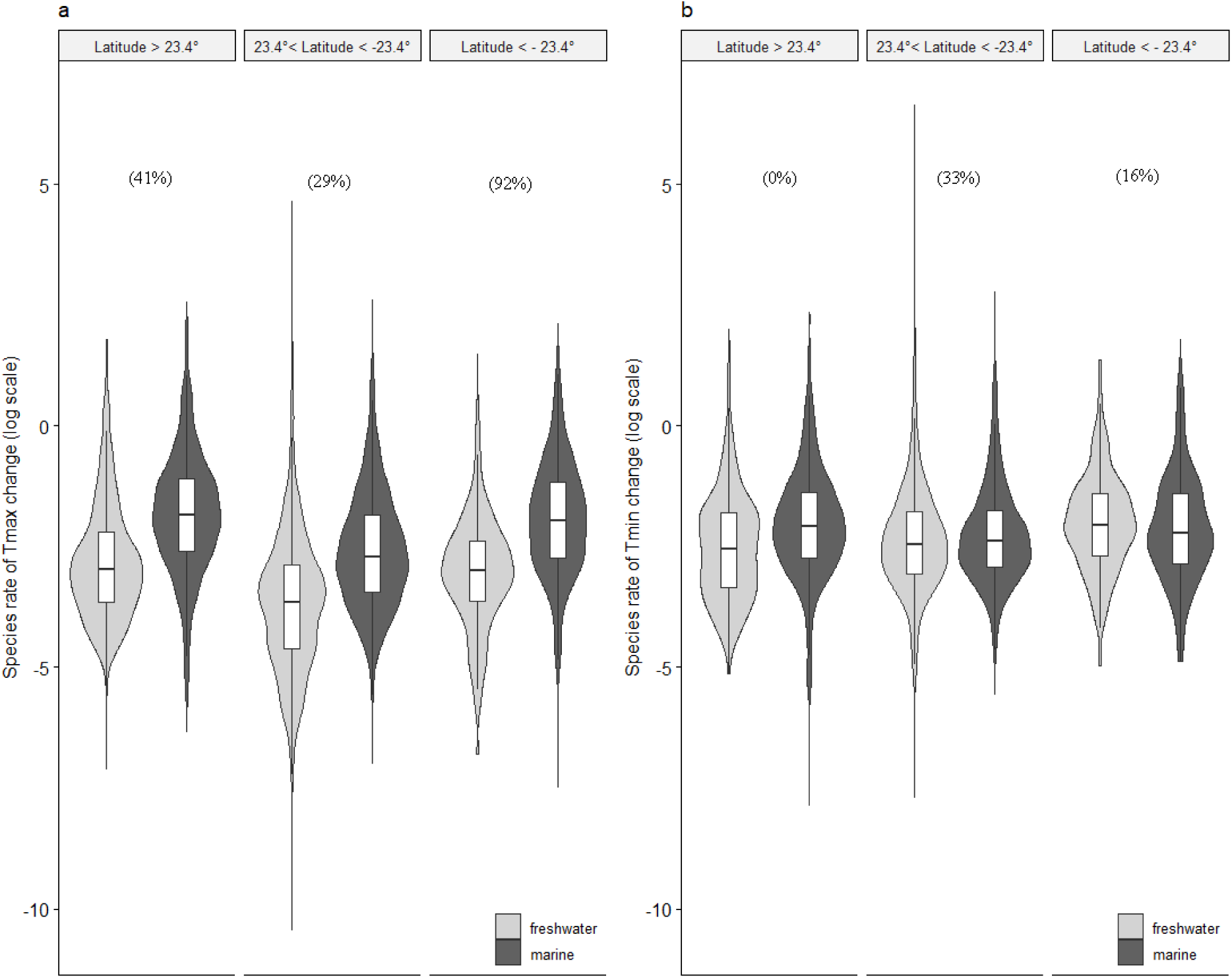
Rates of temperature change for freshwater and marine species according to latitude. The boxes represent the median, the first quartile and the third quartile of species (A) rates of T_max_ and (B) T_min_ change. Violin plots represent sideways density plots of the rate values. 23.4° and −23.4° of latitude are defining the limits between tropical and temperate regions. Pgls performed on the 100 phylogenies showed lower rates of change for freshwater species (Supplementary Table 3). Percentages above the violin plot show the proportion of significant Pgls tests.

Along the latitudinal gradient, the association between latitude and rates of climatic niche change was stronger for freshwater species (Fig. 1, Fig. 3). We propose two possible explanations for the steeper increase of rates of climatic niche change with latitude for freshwater species: first, the relationship between climatic niche breadth and latitude is steeper for freshwater fish (Supplementary Fig. 4). We suggest that this could strongly accelerate climatic niche change for freshwater species at high latitude and offset their overall lower rates of climatic niche change. Indeed, consistently with the literature, we found a positive relationship between rates of temperature change and niche breadth (Fig. 2). Temperate species having wider niches seem to be more prone to climatic niche change^13, 14^. On the contrary, climatic specialization in the tropics may hinder climatic niche change as species with narrow niches might be constrained in their ability to evolve their thermal tolerances or to expand their geographical ranges^14^. Niche breadth showed a steeper increase with latitude for freshwater than for marine species (quadratic effect of latitude on niche breadth average from 100 pgls tests for freshwater species: 0.017 °C/° of latitude and *P*<0.001 against 0.0005 °C/° of latitude and *P* <0.001 for marine species, Supplementary Fig. 4) which might explain the higher increase in rates of temperature change with latitude for freshwater than marine species. Secondly, the relationship between latitude and species distance to their critical thermal maxima seems to be steeper for freshwater species than for marine species^6, 39^. This might favor climatic niche change in the temperate region for freshwater species. However, using the measures of critical thermal maxima (CT_max_) gathered by Comte & Olden^6^, we calculated species thermal safety margin (distance to CT_max_) and found only a weak positive relationship between rates of climatic niche change and thermal safety margin (Supplementary Fig. 7).

By combining an unprecedented dataset of distributional and phylogenetic data for Actinopterygii, we demonstrated that rates of climatic niche change are higher in the temperate regions where climate has been less stable through time. Although we did not account for changes in species geographical distribution^19^ when calculating rates of climatic niche change, we found that the latitudinal pattern still hold for species that have not underwent change in distribution (Appendix 2 and Supplementary Fig. 8). We also showed that the association between latitude and rates of climatic niche change was stronger for freshwater species which also had lower rates of change. We propose that habitat heterogeneity, niche breadth and distance to thermal tolerances explain differences in rates of niche change between freshwater and marine environments. Our results shed a new light on the environmental conditions and species features impacting rates of climatic niche change in aquatic habitats which on the long term, will be key to understand species vulnerability related to climate change.

## Methods

### Phylogenetic reconstruction and dating

We used a three-step hierarchical approach adapted from Chesters^40^ to constrain deep level relationships using taxonomic information and resolve mid and species level relationships from complete mitochondrial DNA and mitochondrial genes: (1) the phylogenetic classification of bony fishes version 4^17^ was used as a backbone topology; (2) Keeping the backbone, we reconstructed a phylogeny from complete mitochondrial DNA for 2,386 species (see Supplementary Methods, Supplementary Fig. 9); (3) the phylogeny from complete mitochondria was used as a constraint to reconstruct the species level phylogeny. Complete mitochondrial data and mitochondrial genes were not combined in the same DNA matrix to avoid poor data overlap thus ensuring homogeneity. Species level relationships were resolved based on two mitochondrial genes classically used for fish barcoding: Cytochrome Oxidase subunit 1 [COX1] and Cytochrome b [CYTB]. All sequences were retrieved from GenBank^41^. GenBank annotations were checked by protein profile identification using MitoPhAST^42^. Alignments were performed using translatorX^43^ and MAFFT v7.222^44^ to ensure conservation of the reading frame. Species names were checked and corrected following the FishBase taxonomy^45^. Rigorous care was taken to remove low quality sequences (see Supplementary Methods). In particular, SATIVA^46^ was used to identify sequences with taxonomic annotation not supported by the tree reconstruction. The final matrix of mitochondrial genes combined 10,576 COX1 and 8,020 CYTB sequences for 12,616 fish species. Tree reconstruction was performed using RAxML v8.2.4^47^. We reconstructed 100 replicate trees to account for the topology uncertainty. Time-calibration of each of the 100 trees was obtained from a secondary calibration approach (see Supplementary Methods) using treePL^48^. We finally provided a species level phylogeny for 12,616 species of Actinopterygii representing around 40% of the known diversity, a larger coverage than the previous phylogenies of Mirande^49^ and Rabosky et al. (when considering the number of species with DNA data)^34^.

### Distribution data

We obtained 10,177,303 present-day geographical occurrences for ‘Actinopterygii’ in the form of geographical coordinates from global and regional databases covering both marine and terrestrial ecosystems (Supplementary Table 5). We then removed duplicated occurrences and unknown species following the FishBase valid taxonomy and synonymies^45^. All species whose range included freshwater habitats were considered as freshwater following Vega & Wiens^27^. We retrieved polygons of estimated species geographical ranges and removed occurrences falling outside the polygons. Polygons were retrieved from the International Union for Conservation of Nature website (IUCN, http://www.iucnredlist.org) (4,762 species) or from Carvajal-Quintero et al.^50^ (2,434 species), both following the same methods. For marine species with no polygons of geographical ranges available (13’185 species), we removed occurrences falling outside the Major Fishing Areas where marine species have been recorded by the Food and Agriculture Organization of the United Nations (FAO, http://www.fao.org/fishery/area/search/en). For freshwater species (4,971 species), we removed occurrences outside the drainage basins where species are known to occur^23^. At this stage, we had compiled a total of 9,042,510 occurrences, 7,928,290 in marine environment and 1,114,220 in freshwaters representing the distribution of 25,812 species. For the following analyses, we only kept species with more than 10 occurrences and selected the species also present in the phylogeny. As later calculation will be performed on each family independently, we extracted the list of monophyletic families in each of the 100 trees. We selected families for which we had distribution data for more than 50% of the species to control the representativeness of our results and families with more than 10 species to ensure model convergence. We obtained a list of 244 families (among the 446 families included in the phylogeny) and kept only the species belonging to these families. This let us with a set of 6,104,127 occurrences representing the distribution of 6,627 species (3,622 marine, 3,005 freshwater species). We obtained for each species an estimate of the mean latitude of occurrence by averaging the latitude of each occurrence point.

### Climatic data

Minimum and maximum temperatures were calculated for each occurrence from different sets of climatic data according to species habitat. We selected two variables instead of performing a Principal Component Analysis on all the climatic variables available (as it was previously done by Kozak & Wiens^3^ and Fisher-Reid et al.^14^) because the relationship between variables can change through time^51^. We used the equation proposed by Punzet et al.^52^ to calculate water temperature on land from the maximum temperature of the warmest month (BIO5) and the minimum temperature of the coldest Month (BIO6) provided by WorldClim v2 (http://www.worldclim.org) (5 arc-minute resolution)^53^. We distinguished marine species living near the surface (reef-associated and pelagic species) or near the bottom (bathydemersal, bathypelagic, benthopelagic and demersal species) as indicated in FishBase^45^ and used, respectively, minimum and maximum sea surface temperature layers or minimum and maximum temperature at maximum depth layers, all downloaded from Bio-ORACLE (Ocean Rasters for Analysis of Climate and Environment, http://www.bio-oracle.org)^54^. When occurrences were covered by both freshwater and marine climatic data (i.e. occurrences for brackish water species), we extracted the temperature using the two types of climatic layers and took the average.

### Rate of temperature niche change

To obtain rates of climatic niche change at the species level, we first calculated for each species an estimate of minimum (T_min_) and maximum (T_max_) temperatures on their current distribution by averaging the temperatures at their occurrences. We decided to investigate both maximum and minimum temperatures as tolerance to heat and tolerance to cold are not always correlated^55^. We modeled temperature change through time using a Brownian motion (BM) process. For each of the 451 monophyletic families from our phylogeny, we fitted a BM model of trait evolution and reconstructed ancestral values at the internal nodes of the tree using the mvMORPH^56^ package in R (functions mvBM and estim). We also calculated at each node the reconstructed niche breadth as the absolute difference between T_min_ and T_max_. We estimated rates of climatic niche change in 3 different ways: (1) rate of temperature niche change, (2) rate of absolute temperature change and (3) rate of niche breadth change. Rates of temperature (T_max_ and T_min_) niche change on each branch were obtained by calculating the difference between the descendant and the ancestral values divided by the branch length. Rates of niche breadth were calculated using the same procedure. Rates of absolute temperature niche change were calculated as the absolute difference between the ancestral and the descendant values divided by the branch length. Each species was assigned to all the branches of the phylogeny joining this species to the root of its family. For each species, we extracted the rates from all the branches it was assigned to and down-weighted each rate by a factor 2 at each step toward the root because the evolutionary history carried by each branch is shared by more and more species when we go back in time (Supplementary Fig. 10). We finally computed the average of these rates, also averaged over the 100 trees of our phylogeny. Rates of temperature and niche breadth change were compared to species mean latitude using linear mixed models in a Bayesian framework (MCMCglmm) while controlling for the sampling of the clade (number of species of the family analysed) and the clade age. Models were run on the 100 phylogenies using the package Multree which incorporates checking of model convergence^57^. The Markov chain Monte Carlo (MCMC) simulations of model parameters were run using an uninformative prior for 1.e^05^ iterations, eliminating the first 100 samples as burn-in, and thinning to every 100^th^ sample. All other comparisons were performed using phylogenetic Generalized least squares model to account for the phylogenetic relatedness between species.

## Supporting information

Details on species included in the phylogeny

Estimated climatic niche

## Supplementary methods

### Phylogenetic reconstruction

Our goal was to provide the most species rich global mega phylogeny of fish while maintaining a consistent backbone and correct branch lengths in order to use it in macro ecological studies. We used a method similar to the one proposed to reconstruct a species-level tree of life for Insects^40^. The method uses a 3 steps hierarchical approach to combine mitochondrial genetic markers with taxonomic information (see Supplementary Fig. 9 for a diagram summarizing the method). We used three species of rays (*Potamotrygon motoro, Pristis clavata* and *Pristis pectinata)* to root the topology. We provide a species level phylogeny for 12,616 species of Actinopterygians representing around 40% of the known diversity of fish. We do not provide a completely automated pipeline but a succession of automated steps allowing an easy update of the phylogeny in order to exploit the rapidly growing DNA databases (Supplementary Fig. 9).

#### Deep level relationships

We used the fourth version of the phylogenetic classification of bony fish^17^ up to the family level to constrain the deeper relationships. This classification is based on the analysis of molecular and genomic data for *c*. 2,000 species, resolving phylogenetic placement for 80% of the bony fish families.

#### Mid level relationships

We retrieved from GenBank^41^ the list of complete mitochondrion available for Actinopterygii using the R package Rentrez^58^. We first removed from this list sequences of hybrids, unverified species and unidentified species by filtering names containing: “_x_”, “_sp_”, “UNVERIFIED”, “sp.”, “spp.”, “spn.”,“_cf_”, “aff.”. We then verified synonymous or misspelled names using the Fishbase dataset^59^ on R and the online list matching tool of catalogue of life^60^. For the remaining list of sequences, we downloaded the mitochondrial protein-coding sequences and sorted them by gene using MitoPhAST^42^. The software checks the validity of Genbank gene annotation through protein profile identification and performs a first alignment which results in 13 Fasta files containing amino acid sequences for the 13 protein coding genes. We used amino acid sequences to avoid dealing with genetic saturation. From these alignments, we only kept one sequence per species. When multiple sequences were available, we selected the Reference Sequence (RefSeq with accession number starting with ‘NC_’) or the longer one. For each gene independently, a second alignment was performed using the most accurate algorithm (L-INS-I) in MAFFT v7.222^44^. Misaligned sequences were visually identified on Geneious 9.0.5^61^ and deleted. We realigned the alignments if necessary. Alignments were trimmed using Gblocks^62^ with the basic options and concatenated to form a supermatrix using FasConcat v1^63^. At this stage, species having sequences with more than 4 missing genes were removed. Using the final alignment, we performed a preliminary tree search without any constraints. We removed species not clustering in the correct order. We finally obtained a supermatrix of 3,475 amino acids for 2’386 species.

#### Species level relationships

We retrieved information on all available mitochondrial sequences belonging to Actinopterygii on Genbank^41^. As in the previous step, we corrected and checked species names with reference databases; verified gene annotation with protein profiling and selected one sequence per gene and per species. From all the mitochondrial protein coding sequences available, COX1 and CYTB were the most abundant as they are usual barcodes for species identification. Although we could have incorporated all available sequences in the matrix to maximize the number of species, we decided to only keep these two barcodes to minimize heterogeneity. As COX1 and CYTB are short and quite conserved barcodes, we kept nucleotide sequences to catch as much variation as possible. For species having a complete mitogenome, the barcode sequences were replaced by the sequences from the mitogenome. Alignment accuracy was increased by aligning sequences independently for each fish order using TranslatorX^43^. Alignments were checked on Geneious 9.0.5^61^ and concatenated per gene using the option mafft-profile in MAFFT v7.222^44^. For each gene, we selected on Geneious^61^ the largest group of sequences which had at least 100 nucleotide positions in common ^64^ and removed sequences with large gaps. Sequences were trimmed using Gblocks^62^ with basic options. We checked whether identical sequences belonged to closely related species and removed them if not. We used SATIVA^46^ to identify species with taxonomic annotations not supported by the tree reconstructed. We performed several iterations (3 times for CYTB, 10 times for COX1) until no more misplaced species was identified. We only provided information on species families and higher taxonomic ranks in order to detect species not falling in the right family. 246 sequences were removed for COX1, 110 for CYTB. The two matrices of barcode sequences were then concatenated, resulting in a matrix of 1,655 nucleotides for 12,616 species (10,576 species for COX1, 8,020 for species CYTB of which 5,980 species had both COX1 and CYTB). Our pipeline allowed the filtering of 362 sequences of COX1 and 180 of CYTB which was a necessary step to build a reliable phylogeny^64^.

#### Final phylogenetic inference

The phylogenetic classification, the mitogenome matrix and DNA barcode matrix were combined in order to obtain the final species-level tree.

(1) The taxonomic classification of the species in the mitogenome matrix was transformed into a taxonomic tree in R (function as.phylo from the APE package^65^). Monophyly was relaxed when families were indicated as “not validated”, “not examined”, “not monophyletic”, “awaiting for formal description” and for orders having an uncertain position within the series (marked as *Incertea sedis*).

(2) Tree search from the mitogenome matrix was performed with the taxonomic tree as a constraint. In order to find the best partitioning and the best model of evolution scheme, we performed a preliminary estimation using PartitionFinder^66^ on a subset of 362 species representing every family. We subsequently used four partitions and the MtREV model of protein evolution. Maximum likelihood tree search was performed using RAxML 8.2.4^47^ with the GAMMA model of rate heterogeneity. The best ML tree obtained containing 2386 species from 362 families did not have enough species coverage to be a sufficient constraint for the final tree search with 12,616 species. Therefore, we artificially added to the tree from complete mitochondrion every species contained in the barcode matrix to ensure their correct position. Whenever possible, species were added to the tree as polytomies (at the basis of a group with no branch length) in the family they belong to. As not all fish families were represented in the mitogenome tree, 656 species were bound at the order and 50 at the series level.

(3) The tree built from complete mitochondrial data with species attached as polytomies was used as a constraint in the final tree search using the matrix of 12,616 barcode sequences. One partition per gene was used under the GTR model. We reconstructed 100 replicate trees to account for the topology uncertainty (bootstrap search using RAxML 8.2.4 with the additional option –k to obtain branch length on bootstrap trees).

#### Time-calibration

Time calibration was performed using a secondary calibration approach. We retrieved the age of the main monophyletic taxonomic groups defined in Betancur-R et al.^17^, from families to megaclasses. These ages were used as priors to time-calibrate individually the 100 replicate trees. For each of the 100 replicate trees, we only selected ages for which the taxonomic group was also monophyletic in the tree. Dating was performed using TreePL^48^. For each tree, we used the best smoothing parameters obtained from a cross-validation procedure. We finally combined the 100 replicate trees into one phylogeny (multiPhylo object).

## Supplementary results

### Appendix 1: Family rates of temperature change

Besides calculating rates of climatic niche change at the species level, we calculated family rates of climatic niche change. For each monophyletic family extracted from the tree, we modeled temperature change along the phylogeny using a BM model of trait evolution. We extracted the evolutionary rate (sigma) as an estimate of the rate of temperature change at the family level. We considered families to be freshwater or marine when they contained respectively more than 75% of freshwater or marine species and labeled families ‘mix’ otherwise. As for rates at the species level, we found that freshwater families have lower rate of T_max_ change than marine families (Supplementary Fig. 6).

### Appendix 2: Species rates of temperature change are lower for tropical than temperate species when considering species that have been restricted to one of these region in the recent past

In our analyses, we assumed species distributions to be fixed through time. However, species distribution likely changed during their evolutionary history and species current position on the latitudinal gradient may not reflect their past position thus skewing the latitudinal pattern we present. We tested whether the difference between rate of climatic niche change in the tropics and the temperate zone still holds when considering only species that have been restricted to one zone since the emergence of their family, temporal scale at which we do our calculation. In order to select species that have been restricted either to the tropics or the temperate region since the emergence of their family, we selected species belonging to families containing only species currently restricted to one of the two regions. First, species were treated as currently restricted to one region when their whole distribution was within this region : minimum and maximum latitude contained in the area (56 temperate and 67 tropical species from 13 families). As this yielded very few species, in a second comparison, species were treated as currently restricted to one region when their mean latitude fell within this region (351 temperate and 942 tropical species from 97 families). Again, we found that for these selected species that may not have underwent shifts in distribution during their evolutionary history, rates of temperature change were lower for tropical than for temperate species (Supplementary Fig. 8).

### Appendix 3: Trends are maintained when removing outlier rates of temperature change

Although, extreme rate values have already been filtered when selecting families with more than 10 species and more than 50% of their species covered, extreme values might still be artifacts coming from unresolved and very short branches in the phylogeny. To ensure that these extreme values do not affect the latitudinal gradient we demonstrate, we tested for the latitudinal gradient while removing the outliers’ values outside the inter-quartile range of rate values. We found again a significant and positive quadratic effect of latitude on rate of temperature change (Supplementary Fig. 11, Supplementary Fig. 12).

### Appendix 4: Robustness of rate of temperature change estimation

To test for the robustness of our rate of temperature change estimation, we compared the rates of temperature change calculated in the main text with those obtained from the last branch leading to the species as done in Quintero & Wiens ^9^ and Comte & Olden^6^. We found that rates calculated with the two methods are highly correlated (Supplementary Fig. 13). We also found that the phylogeny recently published by Rabosky et al.^34^ produced similar rates (Supplementary Fig. 14).

## Supplementary figures

**Supplementary Figure 1:**
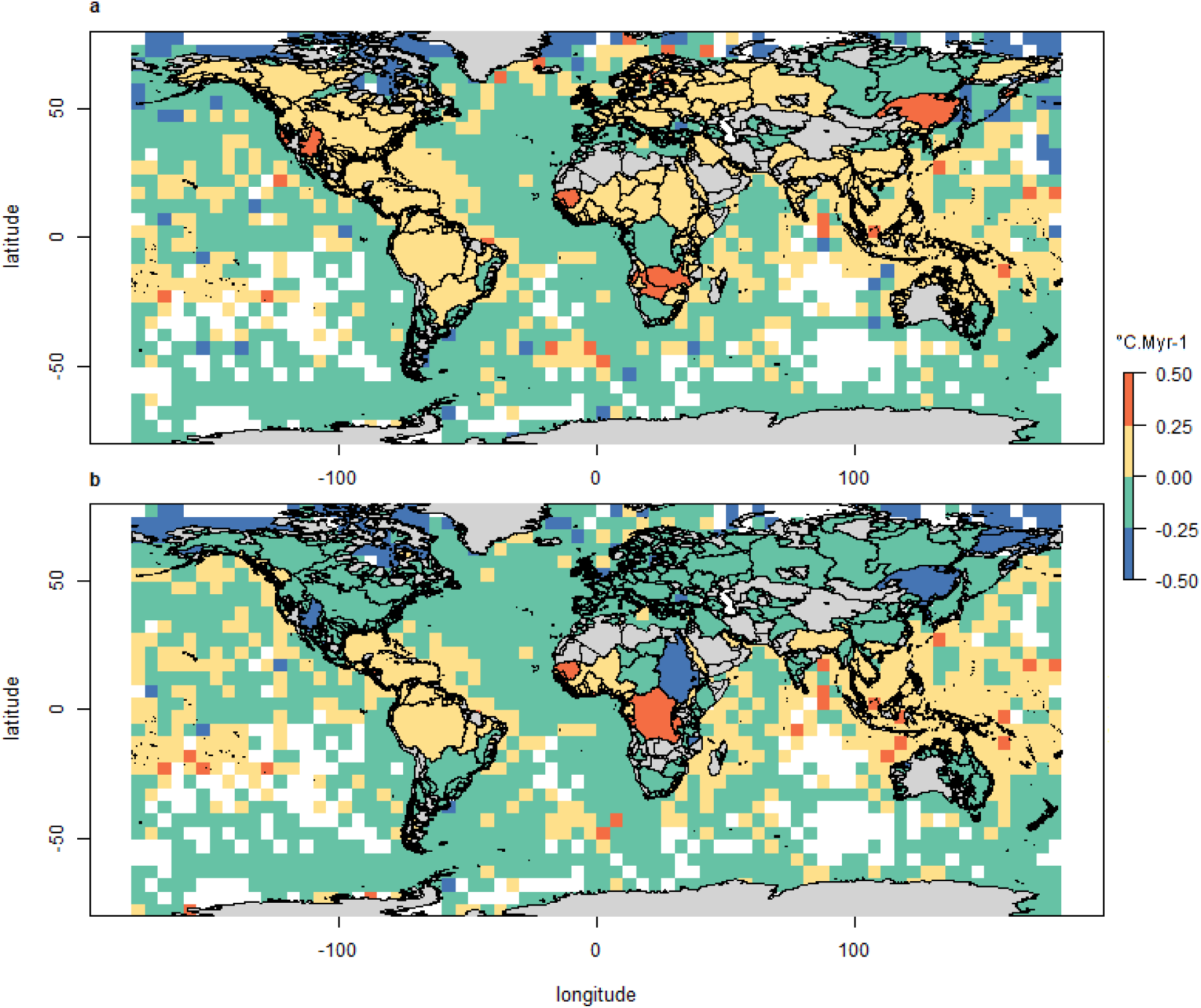
Geographical pattern of rates of non absolute temperature change. Distribution of rate of maximum (T_max_, a) and minimum (T_min_, b) temperature change across the globe (mean rate for species community in each grid cell for marine species and mean rate for fish community in each drainage basin for freshwater species). Rate values were transformed using the square root of the absolute values before putting the sign back. Positive and negative rates mean respectively change toward hotter or colder temperatures.

**Supplementary Figure 2:**
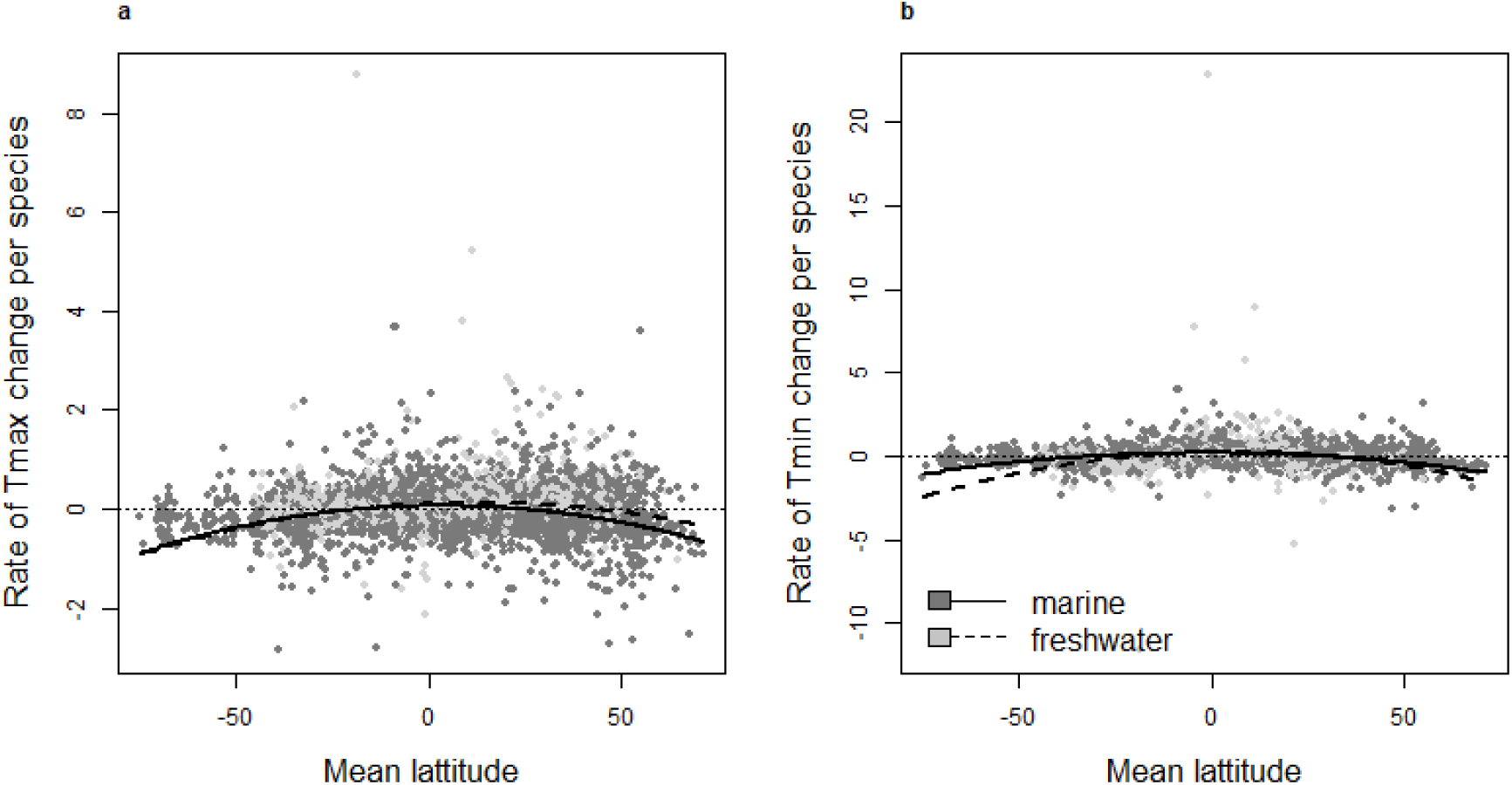
Rates of non absolute T_max_ (a) and T_min_ (b) change according to species mean latitude. Rate values were transformed using the square root of the absolute values before putting the sign back. Positive and negative rates mean respectively changes toward hotter or colder temperatures. Using Bayesian generalized linear mixed-effect models to account for phylogenetic relatedness between species, we found a negative quadratic effect of latitude on rate of temperature change. Thick and dashed lines are built from the median value of the model posterior distribution.

**Supplementary Figure 3:**
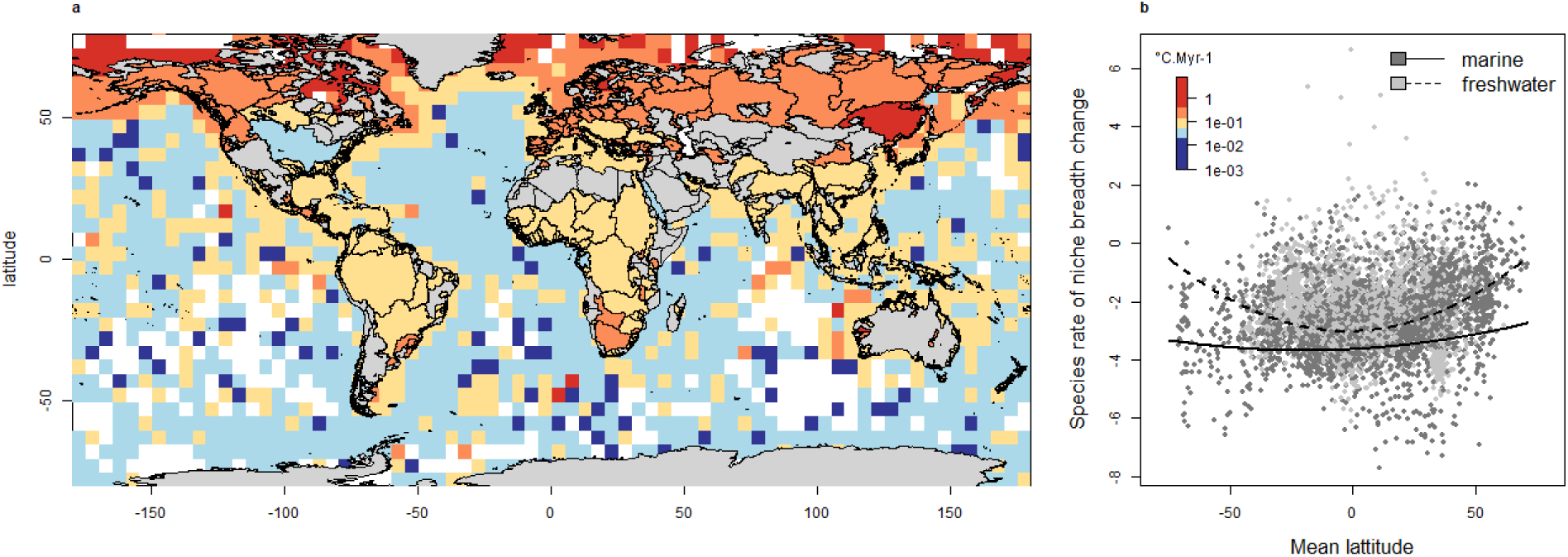
Geographical pattern of rates of niche breadth change. As temperatures were reconstructed for ancestral nodes along the phylogeny, we calculated niche breadth at each ancestral node as the absolute difference between the reconstructed maximum and minimum temperatures. We then calculated rate of absolute niche breadth change for each species as previously done for rate of temperature change. (a) Distribution of rate of absolute niche breadth change (mean rate for the species community in each grid cell for marine species and mean rate for the fish community in each drainage basin for freshwater species). (b) Rate of niche breadth change according to species mean latitude. Using Bayesian mixed-effect models, we found a positive quadratic effect of latitude for both marine and freshwater species. Thick and dashed lines are built from the median value of the model posterior distribution.

**Supplementary Figure 4:**
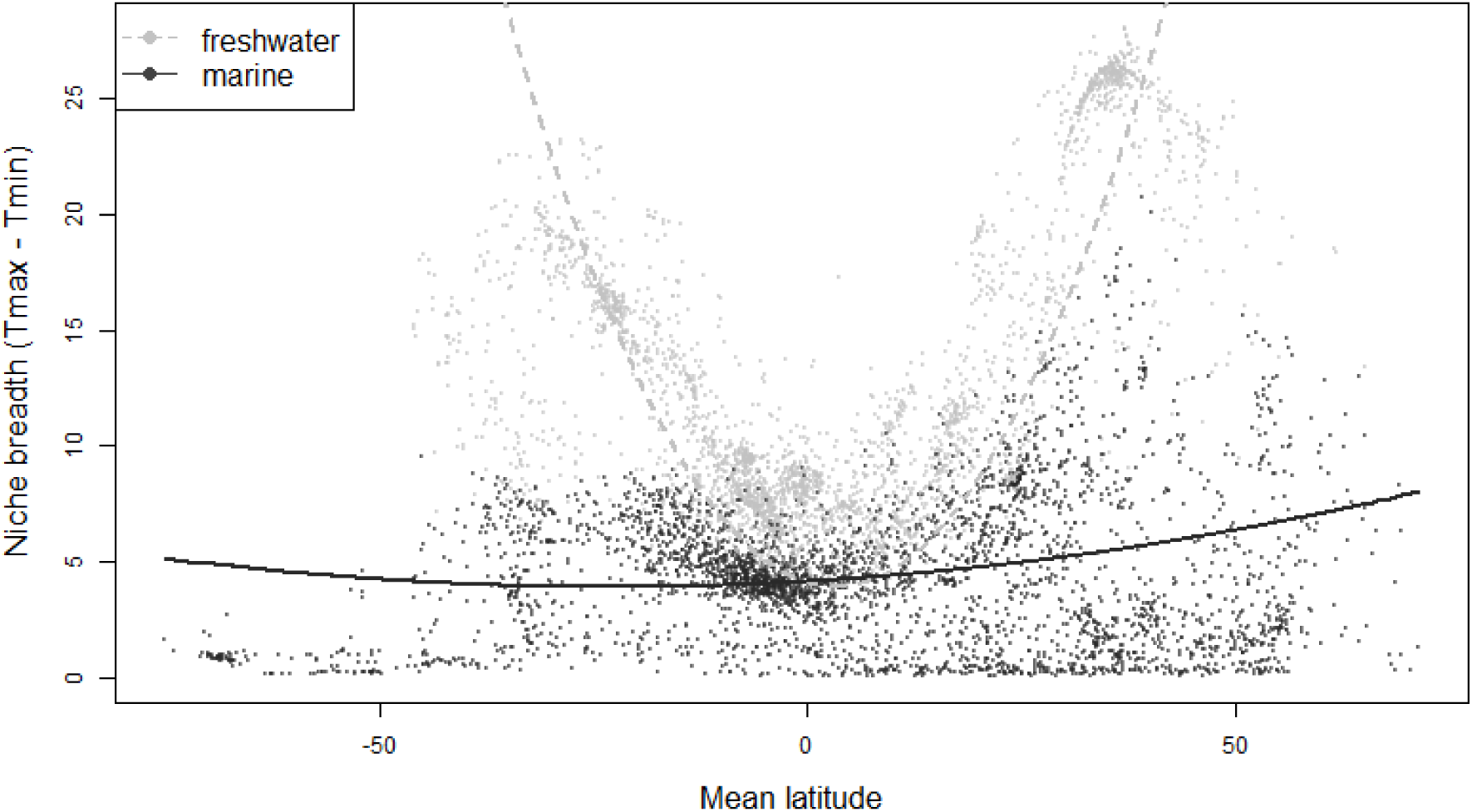
Climatic niche breadth according to species mean latitude. Climatic niche breadth was calculated as the difference between T_max_ and T_min_ across species current distribution. Pgls showed a significant quadratic effect of latitude for both freshwater (quadratic effect of latitude averaged across the 100 pgls tests for freshwater: 0.017, all p-values <0.001, light grey and dashed curve) and marine species (0.0005, all p-values <0.001, dark grey curve).

**Supplementary Figure 5:**
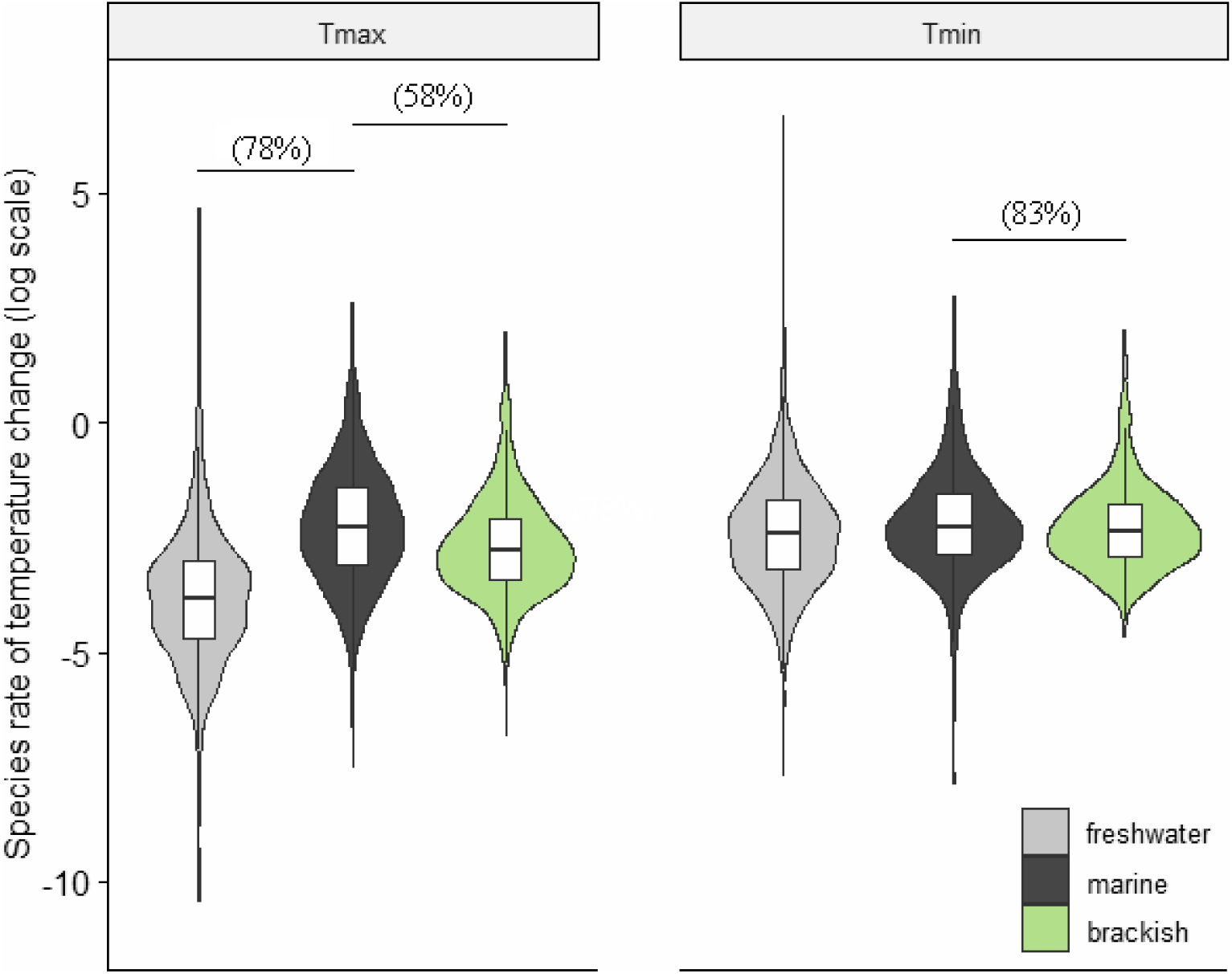
Rate of temperature change for freshwater (light grey), marine (dark grey) species and brackish species (green). The boxes represent the median, the first quartile and the third quartile of the rate of change for maximum (T_max_, left) and minimum (T_min_, right) temperature reconstructed for each species. Violin plots represent sideways density plots of the rate values. Phylogenetic generalized least square regressions were performed on the 100 phylogenies. Percentages above the violin plot show the proportion of significant tests when different from 0. See Table S6 for detailed results.

**Supplementary Figure 6:**
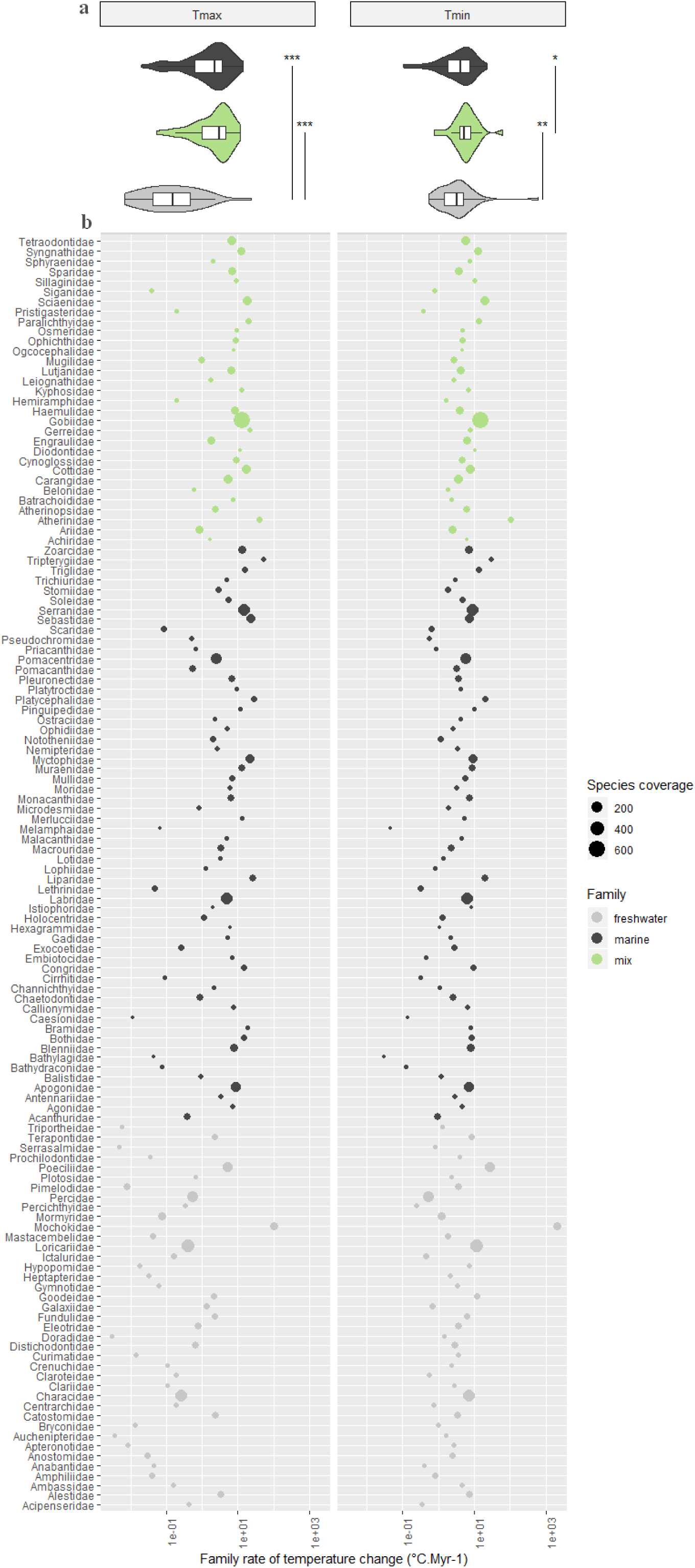
Family rates of temperature change. (a) The boxes represent the median, the first quartile and the third quartile of the family rate of maximum (T_max_, left) and minimum (T_min_, right) temperature change for freshwater (light grey), marine (dark grey) and mixed families (green). Violin plots represent sideways density plots of the rate values. Wilcoxon signed-rank tests showed lower rate of T_max_ change for ‘freshwater’ families than ‘marine’ families (***p < 0.001 for T_max_) and lower rates of temperature change for ‘freshwater’ families than mixed families (***P< 0.001 for T_max_ and **P=0.002 for T_min_). Rates of Tmin change were lower for ‘marine’ families than mixed families (*P=0.048). (b) Rate of temperature change for each family with dot size representing species coverage.

**Supplementary Figure 7:**
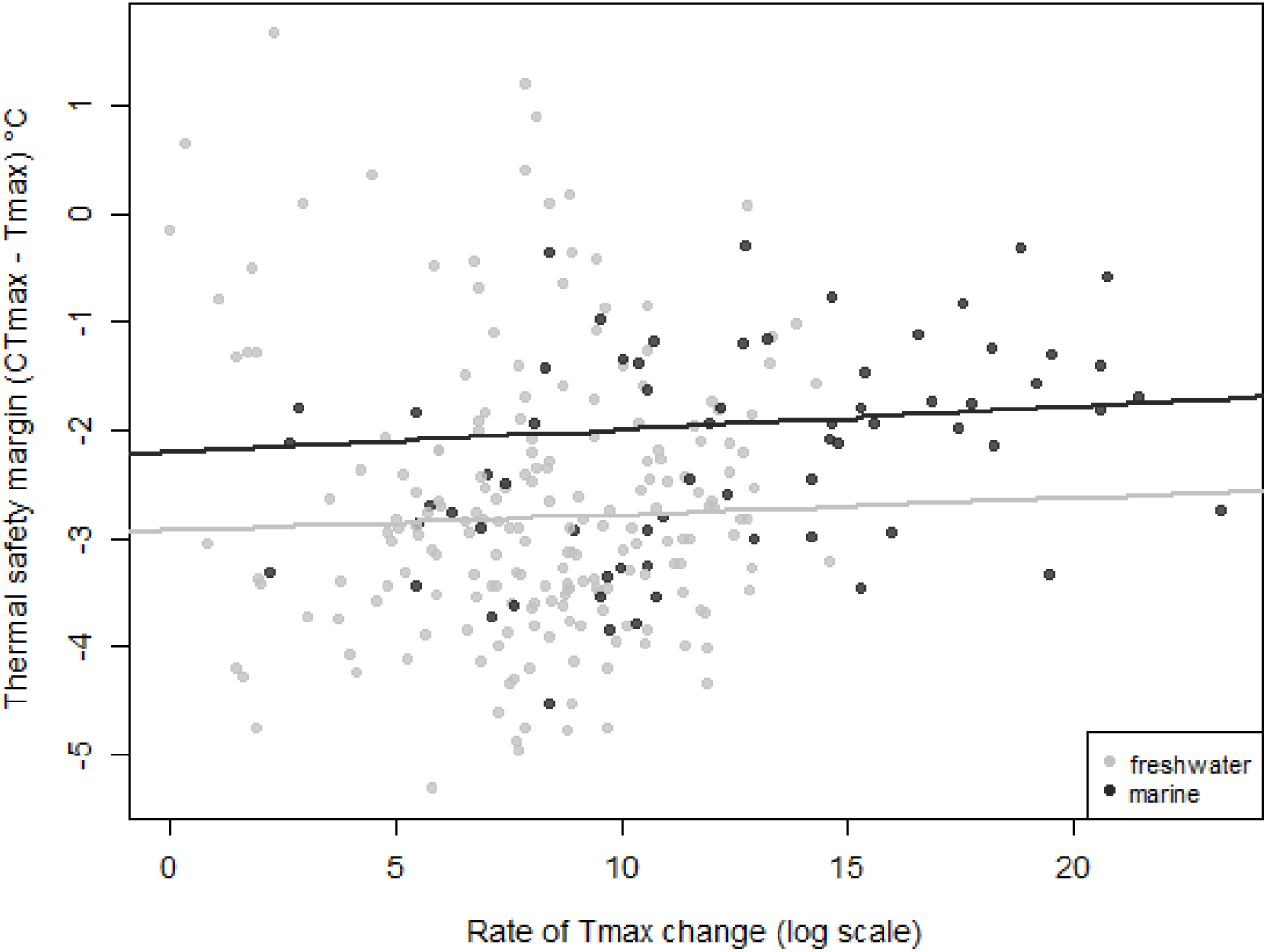
Species rates of T_max_ change according to thermal safety margin (CT_max_-T_max_). We obtained CT_max_ values from Comte & Olden (2017) and took the averaged value for each of the 200 freshwater and 67 marine species. Thermal safety margin was calculated as the difference between CT_max_ and T_max_. Phylogenetic generalized least square regressions were performed on the 100 trees. We found a positive relationship between rate of niche change and thermal safety margin for freshwater (mean effect of thermal safety margin on rates of T_max_ change: 0.015, 11% of significant tests) and marine species (0.021, 6% of significant tests).

**Supplementary Figure 8:**
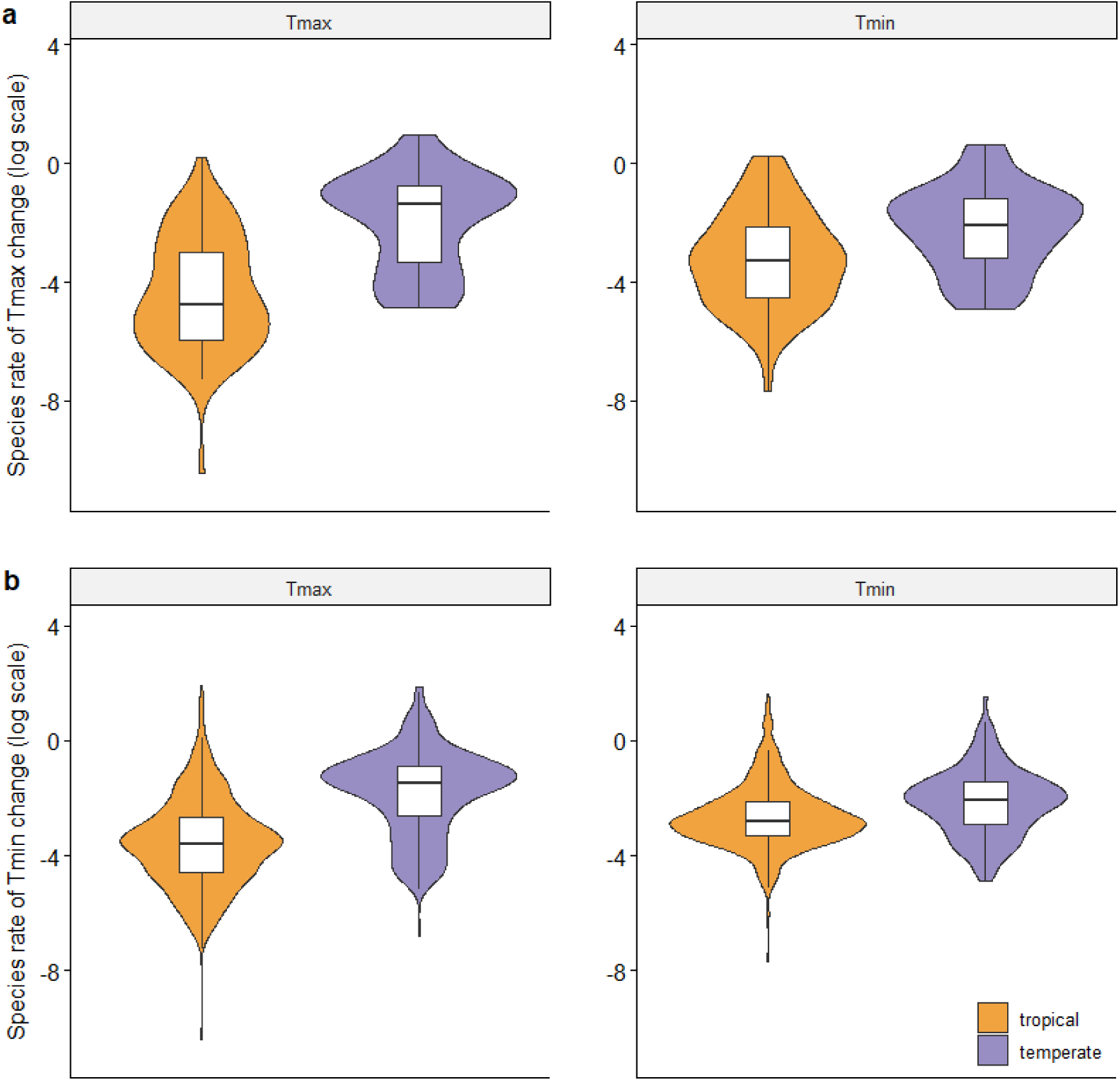
Species rate of T_max_ (left) or T_min_ (right) change according to species latitudinal distribution. We selected species belonging to families containing only species currently restricted to the tropics (latitude −30° to 30°) or the temperate region. (a) Species were treated as restricted to one region when the whole distribution was within this region : minimum and maximum latitude contained in the area (56 temperate and 67 tropical species from 13 families). (b) Species were treated as restricted to one region when their mean latitude fell within this region (351 temperate and 942 tropical species from 97 families). Wilcoxon signed-rank tests showed lower rate of temperature change in the tropical than in the temperate species (P < 0.001 for both T_max_ and T_min_). Pgls did not show a significant difference in rates for tropical or temperate species.

**Supplementary Figure 9:**
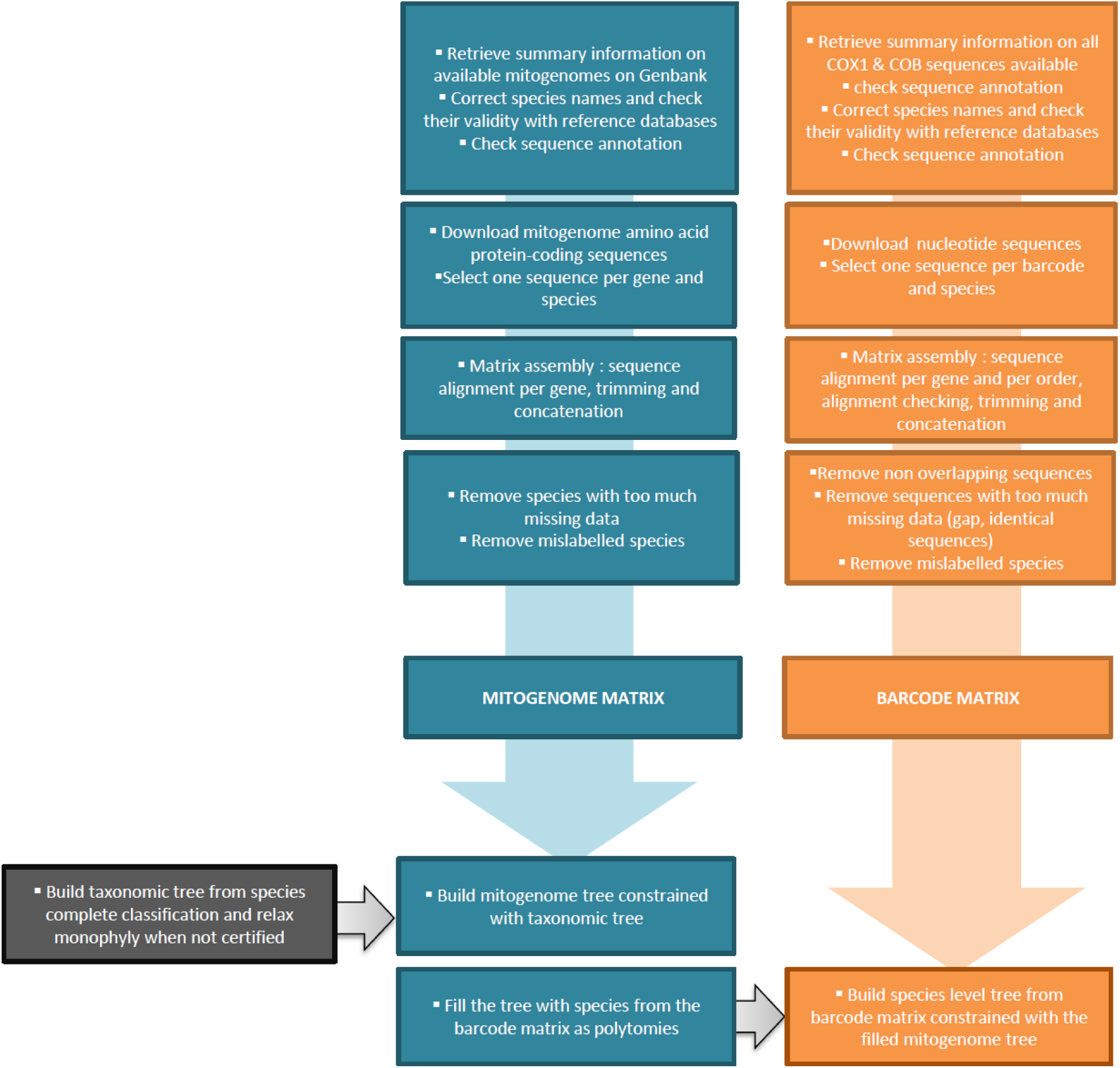
Diagram summarizing the 3 step method used to reconstruct the species-level phylogeny of fish.

**Supplementary Figure 10:**
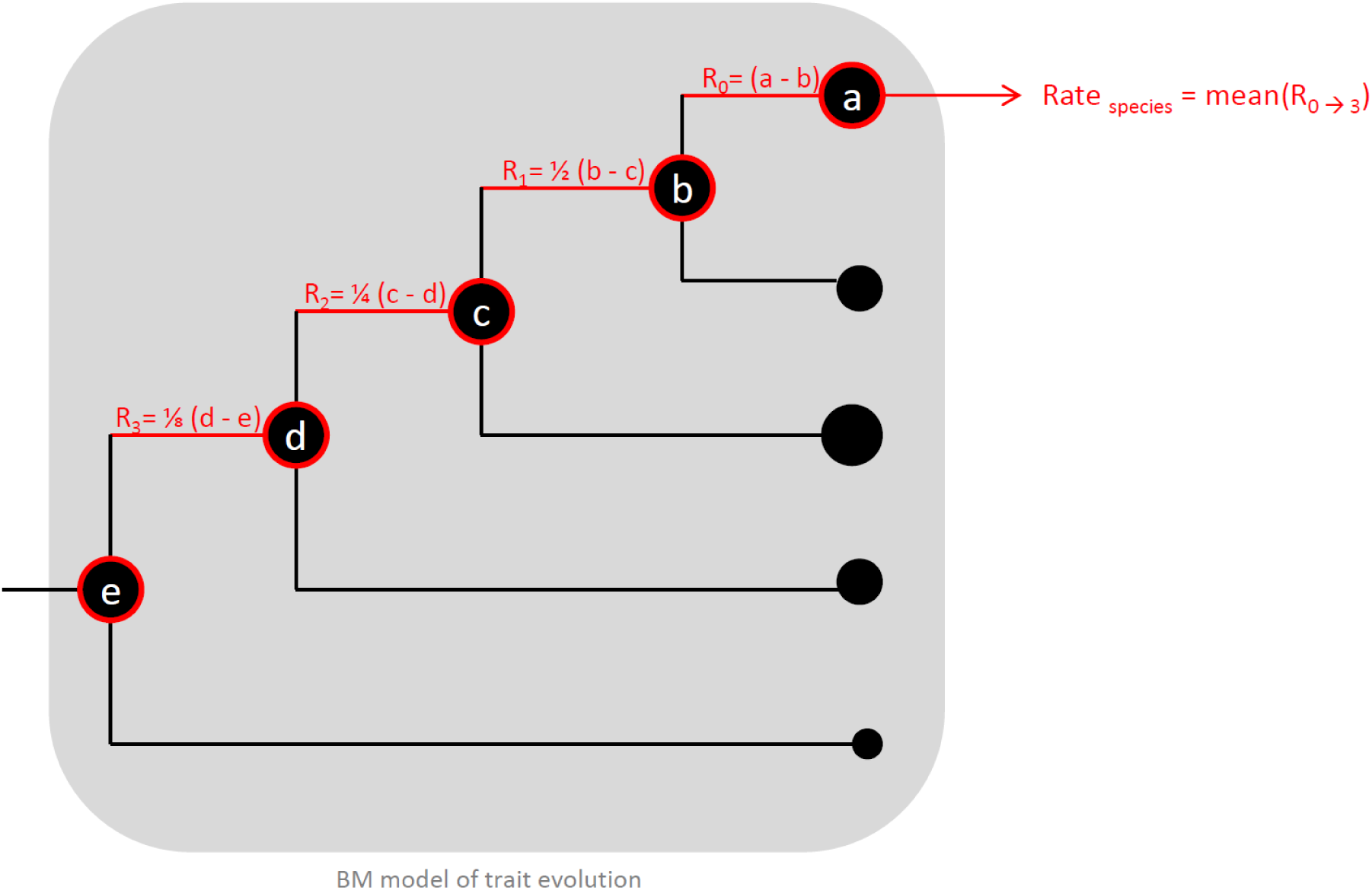
Estimation of species rate of temperature change. The example shows a hypothetic ultrametric tree of a monophyletic family of fish that would have been extracted from the main phylogeny. At the tip of the tree, circles of differing size represent species current climatic niche estimated as the temperature (T_max_ or T_min_) measured on species current distribution. Using a Brownian motion model of evolution, we estimated the ancestral value at each node within the phylogeny (black circles). For each branch, rates were calculated as the difference between the descendant and the ancestor values. For each species, we extracted the rates from all the branches of the phylogeny joining the species tip to the root of its family (red path). Rates on selected branches were down-weighted by a factor 2 at each step toward the root because the evolutionary history carried by each branch is shared by more and more species when we go back in time. We obtained niche change for each species as the average of the down-weighted rates obtained on each branch. This procedure was repeated independently on 100 trees and rates were averaged for each species.

**Supplementary Figure 11:**
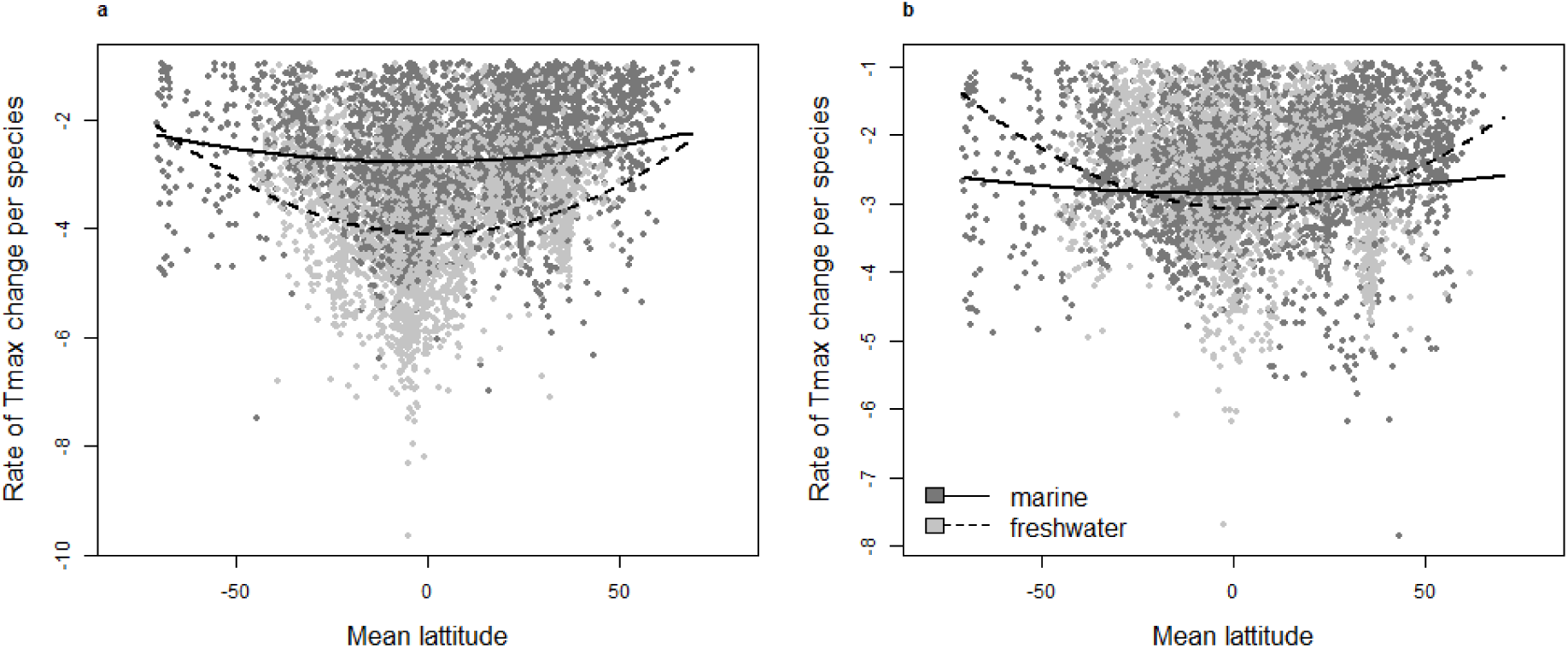
Relationship between rate of T_max_ (a) and T_min_ (b) change and species mean latitude without outlier rate values. Curves are built from the output of generalized linear mixed-effect models relating rate of temperature change to latitude² and accounting for phylogenetic relatedness between species. We found a significant quadratic effect of latitude on rate of temperature change. Thick and dashed lines are built from the median value of the model posterior distribution.

**Supplementary Figure 12:**
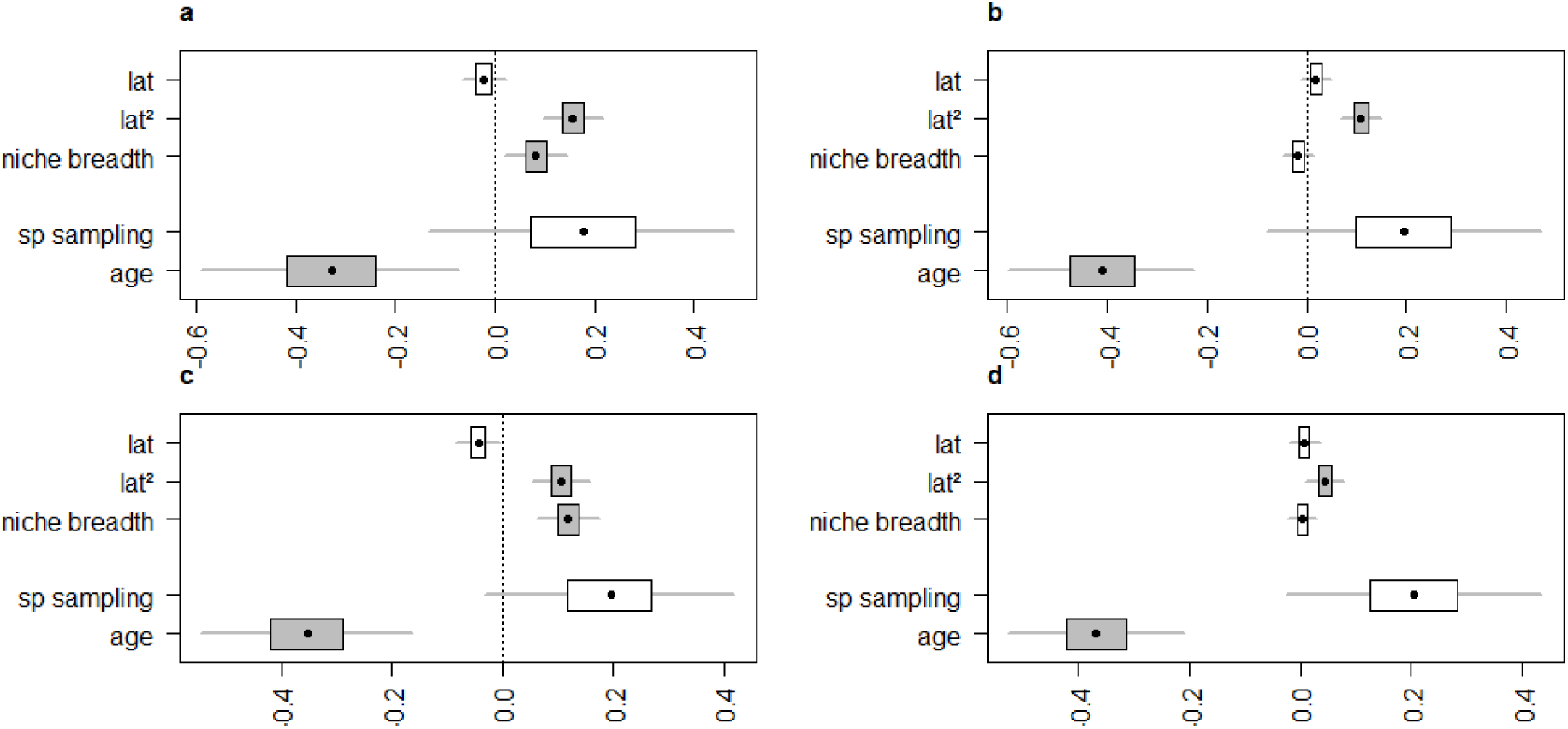
Effect of latitude, niche breadth, species sampling and family age on rates of T_max_ (A,B) and T_min_ (C, D) change for freshwater (A, C) or marine (B, D) species without outlier rate values. We used generalized linear mixed-effect models in a Bayesian framework to account for phylogenetic relatedness between species. The species sampling is the number of species in the family analysed. Models were run on the 100 phylogenies using the R package Multree^57^.

**Supplementary Figure 13:**
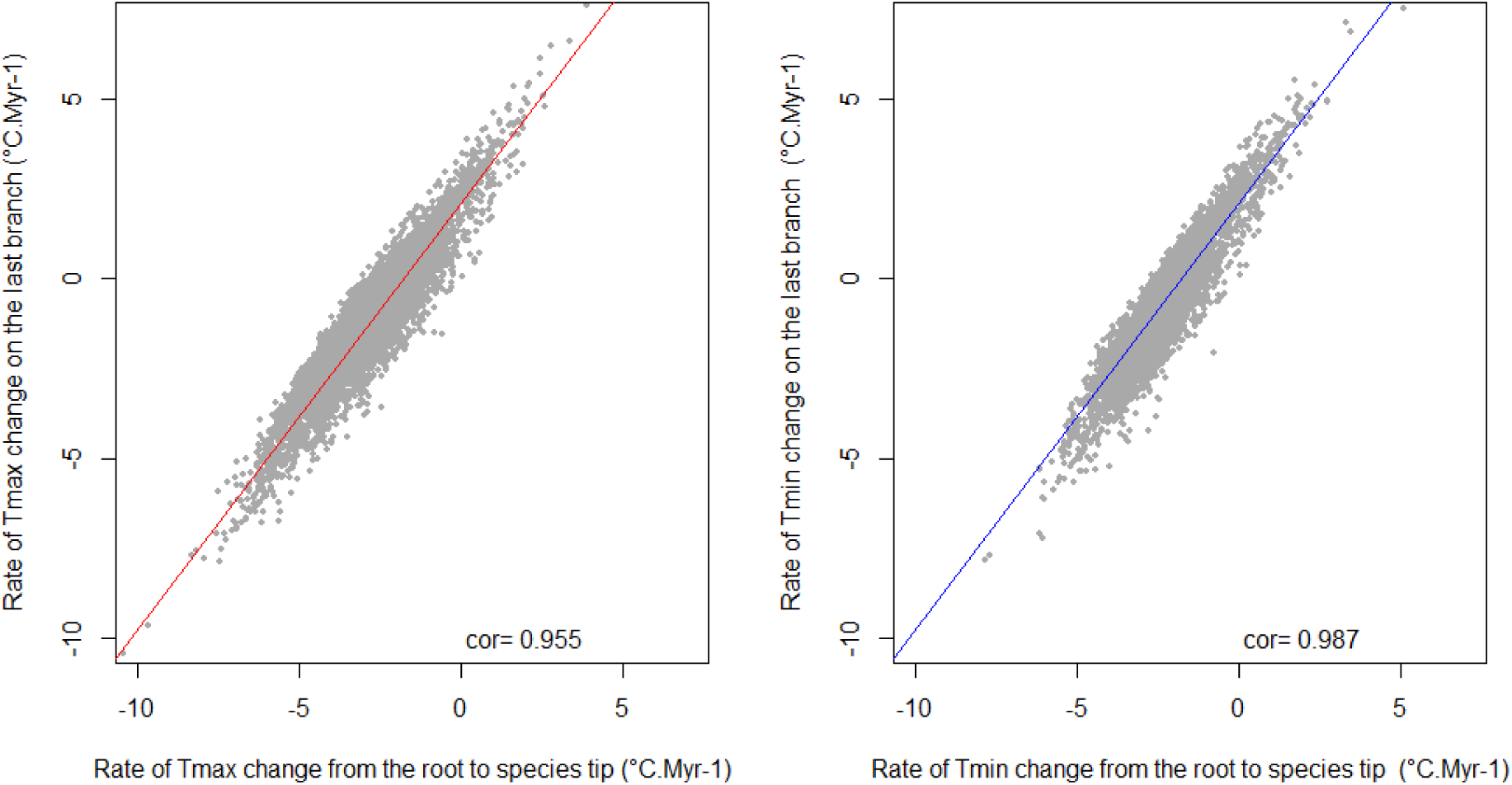
Correlation between rates of maximum (T_max_, left) and minimum (T_min_, right) temperature change used in the main text or calculated on the last branch of the tree (log scale).

**Supplementary Figure 14:**
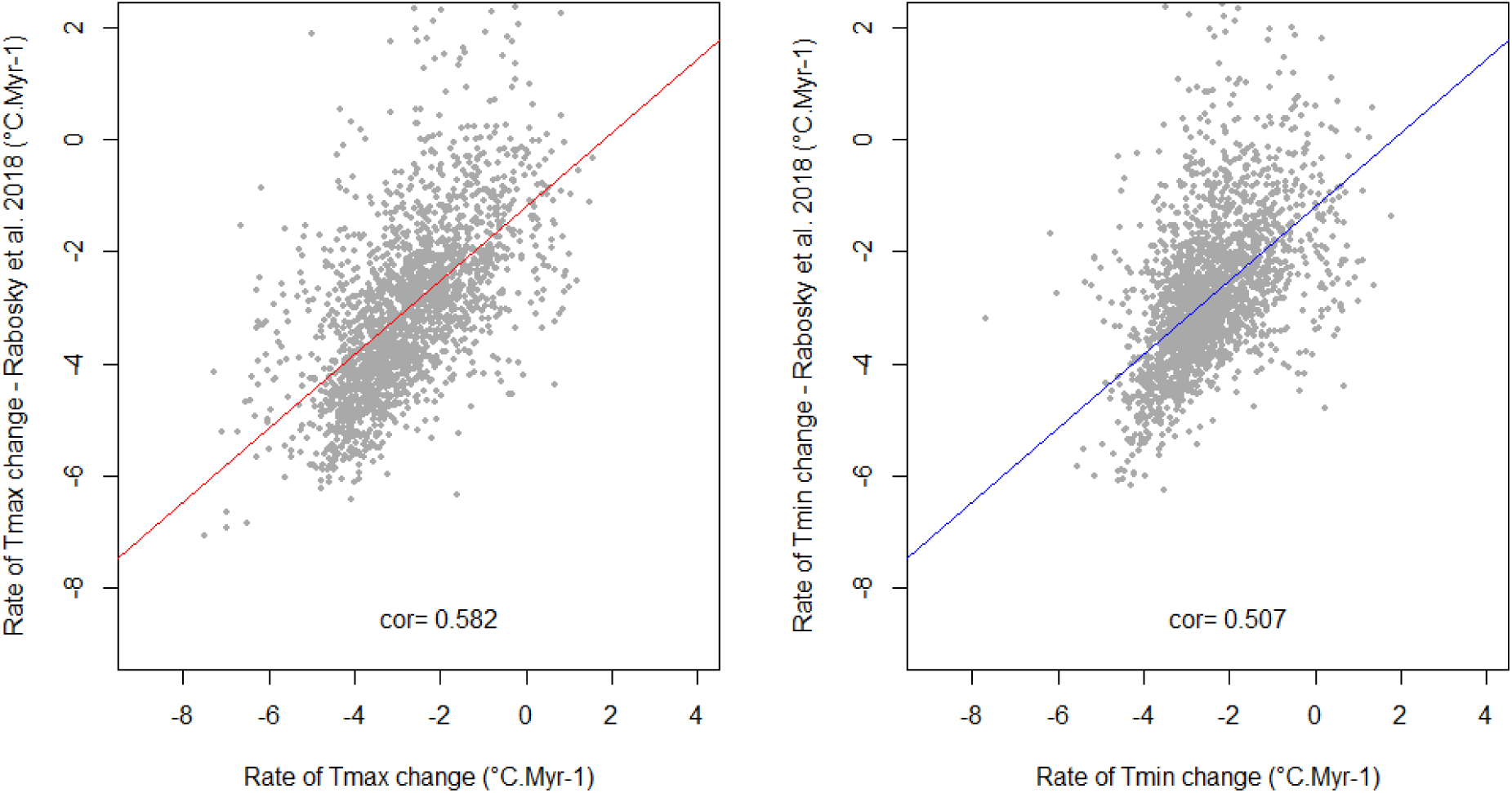
Correlation between rates of maximum (T_max_, left) and minimum (T_min_, right) temperature change as used in the main text or calculated from the phylogeny by Rabosky et al.^34^ (log scale). We followed the same method as in the main text to calculate rates of temperature change using the phylogeny from Rabosky et al.^34^. Here, we compare rates for the 2,114 species representing 82 families that were present in the analyses using the two different phylogenies.

**Supplementary Table 1:** Results of linear models relating rate of temperature change with latitude.

|  |  |  | Estimate | Std.Error | t | pvalue |
| --- | --- | --- | --- | --- | --- | --- |
| **Freshwater** | **T<sub>max</sub>** | Intercept | -3.757 | 0.033 | -113.760 | < 0.001 |
|  |  | latitude | 0.006 | 0.001 | 4.440 | < 0.001 |
|  |  | latitude<sup>2</sup> | 0.001 | 0.000 | 14.780 | < 0.001 |
| **Marine** | **T<sub>max</sub>** | Intercept | -2.566 | 0.026 | -97.872 | < 0.001 |
|  |  | latitude | 0.006 | 0.001 | 7.618 | < 0.001 |
|  |  | latitude<sup>2</sup> | 0.000 | 0.000 | 16.299 | < 0.001 |
| **Freshwater** | **T<sub>min</sub>** | Intercept | -2.376 | 0.028 | -85.503 | < 0.001 |
|  |  | latitude | -0.005 | 0.001 | -4.623 | < 0.001 |
|  |  | latitude<sup>2</sup> | 0.000 | 0.000 | 1.863 | 0.063 |
| **Marine** | **T<sub>min</sub>** | Intercept | -2.300 | 0.024 | -97.701 | < 0.001 |
|  |  | latitude | 0.003 | 0.001 | 4.343 | < 0.001 |
|  |  | latitude<sup>2</sup> | 0.000 | 0.000 | 7.070 | < 0.001 |

**Supplementary Table 2:** Results of bayesian mixed-effect models (MCMCglmm). We tested for the effect of latitude and niche breadth on rates of absolute temperature change (log values) for freshwater or marine species while controlling for family age and species coverage (formula: log(rate)∼latitude+latitude²+species.coverage+niche.breadth+family.age). Variables were scaled to obtain comparable estimates. Median estimated effects and confidence intervals (CI (%)) are provided for each variable.

|  |  |  | median<br>estimate | lower.CI<br>(2.5) | lower.CI<br>(25) | upper.CI<br>(75) | upper.CI<br>(97.5) |
| --- | --- | --- | --- | --- | --- | --- | --- |
| Freshwater | T<sub>max</sub> | intercept | -3.846 | -6.611 | -4.779 | -2.893 | -1.064 |
|  |  | latitude | -0.005 | -0.059 | -0.023 | 0.015 | 0.051 |
|  |  | latitude<sup>2</sup> | 0.129 | 0.055 | 0.103 | 0.152 | 0.200 |
|  |  | species sampling | 0.141 | -0.185 | 0.032 | 0.249 | 0.454 |
|  |  | niche breadth | 0.120 | 0.045 | 0.094 | 0.145 | 0.194 |
|  |  | family age | -0.486 | -0.752 | -0.571 | -0.387 | -0.209 |
|  |  | phylogenetic variance | 7.230 | 5.674 | 6.731 | 7.829 | 11.750 |
|  |  | residual variance | 0.441 | 0.260 | 0.419 | 0.464 | 0.514 |
| Marine | T<sub>max</sub> | intercept | -2.013 | -4.134 | -2.749 | -1.302 | 0.094 |
|  |  | latitude | 0.033 | -0.007 | 0.020 | 0.047 | 0.073 |
|  |  | latitude<sup>2</sup> | 0.131 | 0.077 | 0.112 | 0.148 | 0.182 |
|  |  | species sampling | 0.235 | -0.124 | 0.114 | 0.361 | 0.601 |
|  |  | niche breadth | 0.022 | -0.015 | 0.009 | 0.035 | 0.060 |
|  |  | family age | -0.553 | -0.807 | -0.644 | -0.472 | -0.307 |
|  |  | phylogenetic variance | 7.056 | 5.648 | 6.633 | 7.499 | 8.742 |
|  |  | residual variance | 0.337 | 0.267 | 0.318 | 0.360 | 0.426 |
| Freshwater | T<sub>min</sub> | intercept | -2.807 | -5.177 | -3.606 | -2.002 | -0.427 |
|  |  | latitude | -0.052 | -0.105 | -0.069 | -0.032 | 0.002 |
|  |  | latitude<sup>2</sup> | 0.100 | 0.028 | 0.074 | 0.122 | 0.169 |
|  |  | species sampling | 0.151 | -0.118 | 0.060 | 0.241 | 0.418 |
|  |  | niche breadth | 0.180 | 0.108 | 0.157 | 0.208 | 0.256 |
|  |  | family age | -0.439 | -0.675 | -0.518 | -0.359 | -0.205 |
|  |  | phylogenetic variance | 5.258 | 3.831 | 4.821 | 5.743 | 8.749 |
|  |  | residual variance | 0.499 | 0.366 | 0.475 | 0.519 | 0.575 |
| Marine | T<sub>min</sub> | intercept | -2.237 | -4.165 | -2.906 | -1.573 | -0.289 |
|  |  | latitude | 0.020 | -0.019 | 0.006 | 0.033 | 0.058 |
|  |  | latitude<sup>2</sup> | 0.077 | 0.027 | 0.060 | 0.094 | 0.127 |
|  |  | species sampling | 0.221 | -0.112 | 0.107 | 0.333 | 0.552 |
|  |  | niche breadth | 0.032 | -0.006 | 0.018 | 0.043 | 0.067 |
|  |  | family age | -0.435 | -0.662 | -0.512 | -0.355 | -0.204 |
|  |  | phylogenetic variance | 5.976 | 4.740 | 5.592 | 6.426 | 7.421 |
|  |  | residual variance | 0.344 | 0.278 | 0.324 | 0.365 | 0.420 |

**Supplementary Table 3:**
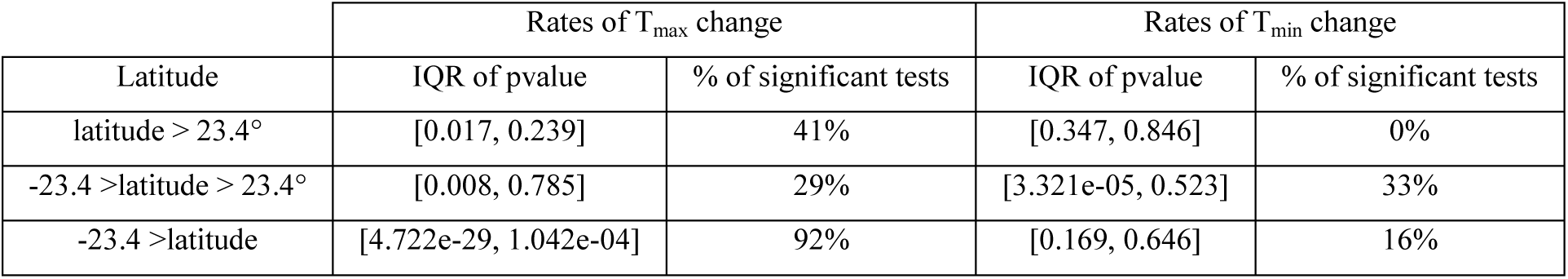
Detailed output of phylogenetic generalized least square (Pgls) regression testing for a difference in rates of temperature change between marine and freshwater species along the latitudinal gradient. Pgls were performed on the 100 phylogenies. The table gives the interquartile range of p-value obtained and the percentage of significant tests as reported in Fig. 3.

**Supplementary Table 4:** Detailed output of phylogenetic generalized least square (Pgls) regression testing for pairwise differences in rates of temperature change between marine, freshwater and brackish species. Pgls were performed on the 100 phylogenies. The table gives the interquartile range of p-value obtained and the percentage of significant tests as reported in Supplementary Fig. 5.

| Comparison | Rates of $T_{\max}$ change | | Rates of $T_{\min}$ change | |
| --- | --- | --- | --- | --- |
|  | IQR of pvalue | % of significant tests | IQR of pvalue | % of significant tests |
| Marine-Freshwater | [0.0003, 0.043] | 78% | [0.514, 0.884] | 0% |
| Marine-Brackish | [0.001, 0.211] | 58% | [5.779e-06, 0.025] | 83% |
| Freshwater-Brackish | [0.550, 0.797] | 0% | [0.473, 0.838] | 0% |

**Supplementary Table 5:** List of databases queried to obtain geographical occurrences for fish species.

| Database | Date | Website | Coverage |
| --- | --- | --- | --- |
| Bold | 02/05/2018 | http://www.boldsystems.org/index.php/ | World |
| Fishnet2 | 02/05/2018 | http://www.fishnet2.net/aboutFishNet.html | World |
| GBIF | 02/05/2018 | https://www.gbif.org/ | World |
| IdigBio | 02/05/2018 | https://www.idigbio.org/ | World |
| Obis | 03/05/2018 | http://iobis.org/ | World |
| FaunAFRI | 03/05/2018 | http://www.poissons-afrique.ird.fr/faunafri/ | Africa |
| Atlas of Life | 02/05/2018 | http://spatial.ala.org.au/webportal/ | Australia |
| Biofresh | 03/05/2018 | http://project.freshwaterbiodiversity.eu/ | Europe |
| SpeciesLink | 02/05/2018 | http://splink.cria.org.br/index?criaLANG=pt | Brazil |
| ICMbio | 02/05/2018 | https://portaldabiodiversidade.icmbio.gov.br/portal/ | Brazil |
| AmazonFISH | 09/04/2018 | https://www.amazon-fish.com/ | Amazon basin |
| Museo Noel Kempf | 05/02/2016 | http://museonoelkempff.org/museo/ | Bolivia |
| PUCRS | 05/02/2016 | http://www.pucrs.br/mct/colecoes/ictiologia/ | Brazil |
| SiBBr | 05/02/2016 | http://www.sibbr.gov.br/ | Brazil |
| Fundacion OGA | 15/06/2010 | https://museoscasso.com.ar/fundacion-oga/ | Argentina |
| NeoDatIII | 30/06/2010 | http://www.mnrj.ufrj.br/search.htm | Brazil |

**Supplementary Table 6:** Relationship between latitude and rates of temperature change. We used bayesian mixed-effect models to estimate and plot (Fig. 1) the relationship between latitude and rates of absolute temperature change (log values) while controlling for the phylogenetic relatedness between species. Median estimated effects and confidence intervals (CI (%)) are provided for each variable.

|  |  |  | median estimate | lower.CI(2.5) | lower.CI(25) | upper.CI(75) | upper.CI(97.5) |
| --- | --- | --- | --- | --- | --- | --- | --- |
| Freshwater | T<sub>max</sub> | intercept | -3.994 | -6.789 | -4.941 | -3.040 | -1.189 |
|  |  | latitude | -0.001 | -0.003 | -0.001 | 0.000 | 0.002 |
|  |  | latitude<sup>2</sup> | 0.000 | 0.000 | 0.000 | 0.000 | 0.000 |
|  |  | phylogenetic variance | 7.353 | 5.815 | 6.847 | 7.959 | 11.996 |
|  |  | residual variance | 0.442 | 0.258 | 0.419 | 0.464 | 0.515 |
| Marine | T<sub>max</sub> | intercept | -2.621 | -4.762 | -3.371 | -1.927 | -0.547 |
|  |  | latitude | 0.001 | 0.000 | 0.001 | 0.002 | 0.003 |
|  |  | latitude<sup>2</sup> | 0.000 | 0.000 | 0.000 | 0.000 | 0.000 |
|  |  | phylogenetic variance | 7.264 | 5.858 | 6.824 | 7.685 | 8.934 |
|  |  | residual variance | 0.334 | 0.263 | 0.314 | 0.356 | 0.421 |
| Freshwater | T<sub>min</sub> | intercept | -2.995 | -5.388 | -3.798 | -2.162 | -0.566 |
|  |  | latitude | -0.003 | -0.005 | -0.004 | -0.002 | 0.000 |
|  |  | latitude<sup>2</sup> | 0.000 | 0.000 | 0.000 | 0.000 | 0.000 |
|  |  | phylogenetic variance | 5.477 | 4.023 | 5.041 | 5.989 | 9.333 |
|  |  | residual variance | 0.496 | 0.351 | 0.474 | 0.520 | 0.576 |
| Marine | T<sub>min</sub> | intercept | -2.746 | -4.682 | -3.411 | -2.086 | -0.812 |
|  |  | latitude | 0.001 | 0.000 | 0.000 | 0.001 | 0.002 |
|  |  | latitude<sup>2</sup> | 0.000 | 0.000 | 0.000 | 0.000 | 0.000 |
|  |  | phylogenetic variance | 6.167 | 4.931 | 5.761 | 6.588 | 7.585 |
|  |  | residual variance | 0.341 | 0.274 | 0.319 | 0.360 | 0.415 |

## Supplementary files

We provide as supplementary files: the list of GenBank accession codes used to build the phylogeny along with species taxonomic classification (accession_phylo_species.xlsx); the estimated temperatures (T_max_ and T_min_) on species distribution and species mean latitude of occurrence (estimated_climatic_niche.xlsx).

## Acknowledgments

We would like to thank Robin Aguilée for his helpful comments on the manuscript along with Fabien Condamine and Jonathan Lenoir for their advice. This work was supported by the EDB Lab which is part of the French Laboratory of Excellence projects ‘LABEX TULIP’ and ‘LABEX CEBA’ (ANR-10-LABX-41, ANR-11-IDEX-0002-02, ANR-10-LABX-25-01). The work was performed using the cluster EDB-Calc (which includes software developed by the Rocks(r) Cluster Group at the San Diego Supercomputer Center, Univ. of California). We are grateful to the AmazonFish project (ERANet-LAC/DCC-0210) for providing species distribution data. LB obtained a PhD funding from the French Ministry of Higher Education, Research and Innovation. J.C.-Q. was funded by Consejo Nacional de Ciencia y Tecnología (CONACYT), Society for Conservation Biology-Latin America & Caribbean Section, and French Biodiversity Agency scholarships.

## Author contributions

LB, GG, JM and JR conceived the study. LB generated the phylogeny with recommendations from GG and JM. JC-Q, CJ and PAT retrieved and filtered distributional data. LB performed the analyses and wrote the manuscript with important contributions from GG, JM and JR. All authors discussed the results and provided inputs on the manuscript.

## References

1. Castro-Insua, A., Gómez-Rodríguez, C., Wiens, J. J. & Baselga, A. Climatic niche divergence drives patterns of diversification and richness among mammal families. Sci. Rep. 8, (2018).

2. Cooney, C. R., Seddon, N. & Tobias, J. A. Widespread correlations between climatic niche evolution and species diversification in birds. J. Anim. Ecol. 85, 869–878 (2016).

3. Kozak, K. H. & Wiens, J. J. Accelerated rates of climatic-niche evolution underlie rapid species diversification. Ecol. Lett. 13, 1378–1389 (2010).

4. Pearman, P. B., Guisan, A., Broennimann, O. & Randin, C. F. Niche dynamics in space and time. Trends Ecol. Evol. 23, 149–158 (2008).

5. Title, P. O. & Burns, K. J. Rates of climatic niche evolution are correlated with species richness in a large and ecologically diverse radiation of songbirds. Ecol. Lett. 18, 433–440 (2015).

6. Comte, L. & Olden, J. D. Climatic vulnerability of the world’s freshwater and marine fishes. Nat. Clim. Change 7, 718 (2017).

7. Cooper, N., Freckleton, R. P. & Jetz, W. Phylogenetic conservatism of environmental niches in mammals. Proc. R. Soc. Lond. B Biol. Sci. 278, 2384–2391 (2011).

8. Jezkova, T. & Wiens, J. J. Rates of change in climatic niches in plant and animal populations are much slower than projected climate change. Proc. R. Soc. B Biol. Sci. 283, 20162104 (2016).

9. Quintero, I. & Wiens, J. J. Rates of projected climate change dramatically exceed past rates of climatic niche evolution among vertebrate species. Ecol. Lett. 16, 1095–1103 (2013).

10. Wiens, J. J. & Graham, C. H. Niche Conservatism: Integrating Evolution, Ecology, and Conservation Biology. Annu. Rev. Ecol. Evol. Syst. 36, 519–539 (2005).

11. Rolland, J. et al. The impact of endothermy on the climatic niche evolution and the distribution of vertebrate diversity. Nat. Ecol. Evol. 2, 459 (2018).

12. Kostikova, A., Litsios, G., Salamin, N. & Pearman, P. B. Linking Life-History Traits, Ecology, and Niche Breadth Evolution in North American Eriogonoids (Polygonaceae). Am. Nat. 182, 760–774 (2013).

13. Smith, S. A. & Beaulieu, J. M. Life history influences rates of climatic niche evolution in flowering plants. Proc. R. Soc. Lond. B Biol. Sci. 276, 4345–4352 (2009).

14. Fisher-Reid, M. C., Kozak, K. H. & Wiens, J. J. How is the rate of climatic-niche evolution related to climatic-niche breadth? Evolution 66, 3836–3851 (2012).

15. Lawson, A. M. & Weir, J. T. Latitudinal gradients in climatic-niche evolution accelerate trait evolution at high latitudes. Ecol. Lett. 17, 1427–1436 (2014).

16. Grosberg, R. K., Vermeij, G. J. & Wainwright, P. C. Biodiversity in water and on land. Curr. Biol. 22, R900–R903 (2012).

17. Betancur-R, R. et al. Phylogenetic classification of bony fishes. BMC Evol. Biol. 17, 162 (2017).

18. Khaliq, I. et al. Global variation in thermal physiology of birds and mammals: evidence for phylogenetic niche conservatism only in the tropics. J. Biogeogr. 42, 2187–2196 (2015).

19. Dynesius, M. & Jansson, R. Evolutionary consequences of changes in species’ geographical distributions driven by Milankovitch climate oscillations. Proc. Natl. Acad. Sci. 97, 9115–9120 (2000).

20. Hua, X. & Wiens, J. J. How Does Climate Influence Speciation? Am. Nat. 182, 1–12 (2013).

21. Schluter, D. Speciation, Ecological Opportunity, and Latitude: (American Society of Naturalists Address). Am. Nat. 187, 1–18 (2016).

22. Weir, J. T. & Price, T. D. Limits to Speciation Inferred from Times to Secondary Sympatry and Ages of Hybridizing Species along a Latitudinal Gradient. Am. Nat. 177, 462–469 (2011).

23. Tedesco, P. A. et al. A global database on freshwater fish species occurrence in drainage basins. Sci. Data 4, 170141 (2017).

24. Hughes, J. M., Schmidt, D. J. & Finn, D. S. Genes in Streams: Using DNA to Understand the Movement of Freshwater Fauna and Their Riverine Habitat. BioScience 59, 573–583 (2009).

25. Nakov, T., Beaulieu, J. M. & Alverson, A. J. Diatoms diversify and turn over faster in freshwater than marine environments. http://biorxiv.org/lookup/doi/10.1101/406165 (2018) doi:10.1101/406165.

26. Seehausen, O. & Wagner, C. E. Speciation in Freshwater Fishes. Annu. Rev. Ecol. Evol. Syst. 45, 621–651 (2014).

27. Vega, G. C. & Wiens, J. J. Why are there so few fish in the sea? Proc R Soc B rspb20120075 (2012).

28. Mora, C. et al. High connectivity among habitats precludes the relationship between dispersal and range size in tropical reef fishes. Ecography 35, 89–96 (2012).

29. Vázquez, D. P. & Stevens, R. D. The Latitudinal Gradient in Niche Breadth: Concepts and Evidence. Am. Nat. 164, E1–E19 (2004).

30. Bush, M. B. & Oliveira, P. E. de. The rise and fall of the Refugial Hypothesis of Amazonian speciation: a paleoecological perspective. Biota Neotropica 6, (2006).

31. Sandel, B. et al. The Influence of Late Quaternary Climate-Change Velocity on Species Endemism. Science 334, 660–664 (2011).

32. Pyron, R. A. & Wiens, J. J. Large-scale phylogenetic analyses reveal the causes of high tropical amphibian diversity. Proc. R. Soc. Lond. B Biol. Sci. 280, 20131622 (2013).

33. Weir, J. T. & Schluter, D. The Latitudinal Gradient in Recent Speciation and Extinction Rates of Birds and Mammals. Science 315, 1574–1576 (2007).

34. Rabosky, D. L. et al. An inverse latitudinal gradient in speciation rate for marine fishes. Nature 559, 392–395 (2018).

35. Schnitzler, J., Graham, C. H., Dormann, C. F., Schiffers, K. & Peter Linder, H. Climatic niche evolution and species diversification in the Cape flora, South Africa. J. Biogeogr. 39, 2201–2211 (2012).

36. Marshall, D. J., Monro, K., Bode, M., Keough, M. J. & Swearer, S. Phenotype– environment mismatches reduce connectivity in the sea. Ecol. Lett. 13, 128–140 (2010).

37. Palumbi, S. R. Marine speciation on a small planet. Trends Ecol. Evol. 7, 114–118 (1992).

38. Bowen, B. W., Rocha, L. A., Toonen, R. J. & Karl, S. A. The origins of tropical marine biodiversity. Trends Ecol. Evol. 28, 359–366 (2013).

39. Pinsky, M. L., Eikeset, A. M., McCauley, D. J., Payne, J. L. & Sunday, J. M. Greater vulnerability to warming of marine versus terrestrial ectotherms. Nature (2019) doi:10.1038/s41586-019-1132-4.

40. Chesters, D. Construction of a Species-Level Tree-of-Life for the Insects and Utility in Taxonomic Profiling. Syst. Biol. syw099 (2016).

41. Benson, D. A. et al. GenBank. Nucleic Acids Res. 41, D36–D42 (2012).

42. Tan, M. H., Gan, H. M., Schultz, M. B. & Austin, C. M. MitoPhAST, a new automated mitogenomic phylogeny tool in the post-genomic era with a case study of 89 decapod mitogenomes including eight new freshwater crayfish mitogenomes. Mol. Phylogenet. Evol. 85, 180–188 (2015).

43. Abascal, F., Zardoya, R. & Telford, M. J. TranslatorX: multiple alignment of nucleotide sequences guided by amino acid translations. Nucleic Acids Res. gkq291 (2010).

44. Katoh, K. & Standley, D. M. MAFFT multiple sequence alignment software version 7: improvements in performance and usability. Mol. Biol. Evol. 30, 772–780 (2013).

45. Froese, R. and D. Pauly. Editors. 2019. FishBase. World Wide Web electronic publication. www.fishbase.org, version (02/2019).

46. Kozlov, A. M., Zhang, J., Yilmaz, P., Glöckner, F. O. & Stamatakis, A. SATIVA : Phylogeny-aware identification and correction of taxonomically mislabeled sequences. Nucleic Acids Res. 44, 5022–5033 (2016).

47. Stamatakis, A. RAxML version 8: a tool for phylogenetic analysis and post-analysis of large phylogenies. Bioinformatics 30, 1312–1313 (2014).

48. Smith, S. A. & O’Meara, B. C. treePL: divergence time estimation using penalized likelihood for large phylogenies. Bioinformatics 28, 2689–2690 (2012).

49. Mirande, J. M. Combined phylogeny of ray-finned fishes (Actinopterygii) and the use of morphological characters in large-scale analyses. Cladistics 33, 333–350 (2017).

50. Carvajal-Quintero, J. et al. Drainage network position and historical connectivity explain global patterns in freshwater fishes’ range size. Proc. Natl. Acad. Sci. 116, 13434–13439 (2019).

51. Uyeda, J. C., Caetano, D. S. & Pennell, M. W. Comparative Analysis of Principal Components Can be Misleading. Syst. Biol. 64, 677–689 (2015).

52. Punzet, M., Voß, F., Voß, A., Kynast, E. & Bärlund, I. A Global Approach to Assess the Potential Impact of Climate Change on Stream Water Temperatures and Related In-Stream First-Order Decay Rates. J. Hydrometeorol. 13, 1052–1065 (2012).

53. Hijmans, R. J., Cameron, S. E., Parra, J. L., Jones, P. G. & Jarvis, A. Very high resolution interpolated climate surfaces for global land areas. Int. J. Climatol. 25, 1965–1978 (2005).

54. Tyberghein, L. et al. Bio-ORACLE: a global environmental dataset for marine species distribution modelling: Bio-ORACLE marine environmental data rasters. Glob. Ecol. Biogeogr. 21, 272–281 (2012).

55. Araújo, M. B. et al. Heat freezes niche evolution. Ecol. Lett. 16, 1206–1219 (2013).

56. Clavel, J., Escarguel, G. & Merceron, G. mv MORPH : an R package for fitting multivariate evolutionary models to morphometric data. Methods Ecol. Evol. 6, 1311–1319 (2015).

57. Healy, K. et al. Ecology and mode-of-life explain lifespan variation in birds and mammals. Proc. R. Soc. B Biol. Sci. 281, 20140298–20140298 (2014).

58. Winter, D. J. rentrez: An R package for the NCBI eUtils API. (2017).

59. Boettiger, C., Lang, D. T. & Wainwright, P. C. rfishbase: exploring, manipulating and visualizing FishBase data from R. J. Fish Biol. 81, 2030–2039 (2012).

60. Roskov Y., Ower G., Orrell T., Nicolson D., Bailly N., Kirk P.M., Bourgoin T., DeWalt R.E., Decock W., Nieukerken E. van, Zarucchi J., Penev L., eds. (2019). Species 2000 & ITIS Catalogue of Life, 2019 Annual Checklist. Digital resource at www.catalogueoflife.org/annual-checklist/2019. Species 2000: Naturalis, Leiden, the Netherlands. ISSN 2405-884X.

61. Kearse, M. et al. Geneious Basic: An integrated and extendable desktop software platform for the organization and analysis of sequence data. Bioinformatics 28, 1647–1649 (2012).

62. Castresana, J. Selection of conserved blocks from multiple alignments for their use in phylogenetic analysis. Mol. Biol. Evol. 17, 540–552 (2000).

63. Kück, P. & Meusemann, K. FASconCAT: convenient handling of data matrices. Mol. Phylogenet. Evol. 56, 1115–1118 (2010).

64. Peters, R. S. et al. The taming of an impossible child: a standardized all-in approach to the phylogeny of Hymenoptera using public database sequences. BMC Biol. 9, 55 (2011).

65. Paradis, E., Claude, J. & Strimmer, K. APE: analyses of phylogenetics and evolution in R language. Bioinformatics 20, 289–290 (2004).

66. Lanfear, R., Calcott, B., Ho, S. Y. & Guindon, S. PartitionFinder: combined selection of partitioning schemes and substitution models for phylogenetic analyses. Mol. Biol. Evol. 29, 1695–1701 (2012).

